# From musk to body odor: decoding olfaction through genetic variation

**DOI:** 10.1101/2021.04.27.441177

**Authors:** Bingjie Li, Marissa L. Kamarck, Qianqian Peng, Fei-Ling Lim, Andreas Keller, Monique A.M. Smeets, Joel D. Mainland, Sijia Wang

## Abstract

The olfactory system combines input from multiple receptor types to represent odor information, but there are few explicit examples relating olfactory receptor (OR) activity patterns to odor perception. To uncover these relationships, we performed genome-wide scans on odor-perception phenotypes for ten odors in 1003 Han Chinese and validated results for six of these odors in an ethnically diverse population (n=364). In both populations, we replicated three previously reported associations (*β*-ionone/OR5A, androstenone/OR7D4, cis-3-hexen-1-ol/OR2J3 LD-band), suggesting that olfactory phenotype/genotype studies are robust across populations. Two novel associations between an OR and odor perception contribute to our understanding of olfactory coding. First, we found a SNP in OR51B2 that associated with trans-3-methyl-2-hexenoic acid, a key component of human underarm odor. Second, we found two linked SNPs associated with the musk Galaxolide in a novel musk receptor, OR4D6, which is also the first OR shown to drive specific anosmia to a musk compound. We also found that the derived alleles of the SNPs reportedly associated with odor perception tend to reduce odor intensity, supporting the hypothesis that the primate olfactory gene repertoire has degenerated over time. This study provides information about coding for human body odor, and gives us insight into broader mechanisms of olfactory coding, such as how differential OR activation can converge on a similar percept.

---

Every individual experiences smell in their own unique way – variation in odor perception can range from specific anosmias, where an individual completely lacks the ability to perceive a particular odorous compound, to differences in individual experience of quality, pleasantness, and/ or intensity of odors(1). Comparing this perceptual variability with genetic variability allows us to identify the role of single odorant receptors in the perceptual code. Progress in sequencing technology and increased access to previously genotyped cohorts has enhanced our ability to uncover the genetic components underlying differences in odor perception.

Olfactory receptors (ORs), the family of proteins responsible for detection of odor compounds, have a high level of genetic variation relative to other proteins(2–4). Of the 800 olfactory receptor genes, only about 400 are intact, and, on average, approximately 30% of OR alleles will differ functionally between two people(5). Even within the set of intact genes, a genetic variant can alter function of a single OR and thereby alter perception of an odor. To date, there are 15 cases where perceptual variability of an odor correlated with a genetic variant in a receptor that responds to the odor in a cell-based assay(5–12).

Here, we utilize the same strategy of correlating perceptual and genetic variation, but with three improvements: *1*. Using a larger population to increase power, *2*. Conducting genetic analysis in a unique and homogenous population (Han Chinese), as opposed to previous studies that have been largely conducted in Western (majority Caucasian) populations, and *3*. Validating the results using an independent population and different methodology, demonstrating the robustness of the finding.

In this study, we tested a Han Chinese population (n=1003) alongside a smaller validation cohort (n=364) of a Western population, using odors that have unexplained variability in perception – Galaxolide, trans-3-methyl-2-hexenoic acid (3M2H), and aldehydes – as well as a set of odors with previously described associations between perceptual variability and genetic variants.

## Galaxolide: A Musk Compound

The olfactory literature contains a number of examples of compounds with very different structures but similar odors(13). The perceptual category of musks is perhaps the most striking example. Compounds in five different musk structural classes – macrocyclic, polycyclic, nitro, steroid-type, and straight-chain (alicyclic) – all have a similar perceptual quality described as sweet, warm, and powdery(14). The simplest explanation is that all musk structures activate one receptor or one common subset of receptors that in turn encodes the perceptual “musk” quality; however, evidence suggests coding of this percept may be more complex. Individuals can have specific anosmias to one or some, but not all musks(15, 16), suggesting that there is not a single common coding mechanism.

In this study, we examined Galaxolide, a musk compound with a characterized specific anosmia(15, 16). Galaxolide does not activate OR5AN1, which was shown to be critical for the perception of other musk compounds in mice(17). The structural variance and common percept amongst musk compounds allows us to examine different coding mechanisms that are central to our understanding of how receptor activation relates to odor perception.

## 3M2H: A Body Odor Contributor

All mammals use chemosensation as a means of intra-species communication, but the mechanism of chemosensory communication amongst humans is largely unknown. The growing evidence for chemical communication between humans suggests that body odor is of particular importance, as it may be processed differently in the brain than other odors(19) and may influence various social behaviors including kinship recognition, mate selection(20), and fear priming(21). Although 3M2H is only one of ∼120 compounds(22) that comprise body odor, it is an “impact odor”, meaning that it carries the characteristic scent of body odor(23). Furthermore, almost 25% of the population has a specific anosmia to 3 M2H(23–26), but this anosmia has not been connected to any olfactory receptor. Identifying receptors responsible for perception of 3M2H and body odor may have implications for social communication, malodor prevention, and receptor coding mechanisms for conspecific odors.

## Replicating Odor Associations

Previous publications have implicated OR genetic variation in perception of specific odors. To examine if these associations are robust and consistent across populations, we measured responses to *β*-ionone(9), androstenone(6, 10), cis-3-hexen-1-ol(8–10, 27), and caproic acid(10).

## Testing Aldehydes in Different Populations

Aldehydes have been shown to vary perceptually across demographic groups such that self-reported Asian populations rate aldehydes as more intense than Caucasian populations(28), but no specific genetic variants or receptors have been implicated. To assess the genetic underpinnings of aldehyde preferences in the Han Chinese population, we tested two monomolecular aldehydes: decyl aldehyde(28) and galbanum oxathiane alongside two fragrance mixtures used in home care products: MixA, which has high levels of aldehydes and is relatively unpopular in Asia, and MixB which has low levels of aldehydes and is popular in Asia.

## Results

To examine genetic variants related to differences in odor perception, we examined how genetic variation correlated with olfactory phenotypes in two cohorts. The discovery cohort consisted of 1003 (370 male) Han Chinese participants, and the validation cohort consisted of 357 (161 male) participants collected in New York City. Participants from both cohorts rated intensity and pleasantness of all odors on a 100-point scale. The discovery cohort had 10 odors presented at a single concentration. Most participants performed each olfactory rating task once, but for each odor a set of 100 participants rated the odor twice throughout the session (test-retest r=0.75). The validation cohort tested 6 of the 10 odors in the discovery cohort, some of which were presented at two concentrations (high/low). Each participant rated all odors twice throughout the session (r=0.69) (Supplementary Fig. 1).

In both cohorts, we normalized participant ratings by ranking across odors by intensity and pleasantness. For the discovery cohort, we performed genome wide association analysis for 20 olfactory phenotypes (Supplementary Data 1). We identified novel genetic variants (Fig. 1) associated with the intensity rankings of Galaxolide and trans-3-methyl-2-hexenoic acid (3M2H) that explain 13.26% and 4.13% of the phenotype variance, respectively. We can compare this to the maximum expected values provided by heritability analysis, which estimates 33% (Galaxolide) and 24% (3M2H) of phenotypic variance is caused by genetic variation (Supplementary Table 1). In addition, we replicated published associations for *β*-ionone, androstenone, and cis-3-hexen-1-ol (Fig. 1). The validation cohort replicated novel associations identified in the discovery study (Galaxolide, 3M2H) as well as published associations (*β*-ionone, androstenone, cis-3-hexen-1-ol). n both cohorts, all the genetic variants that were significantly associated with pleasantness were also significantly associated with intensity perception.

**Fig. 1.**
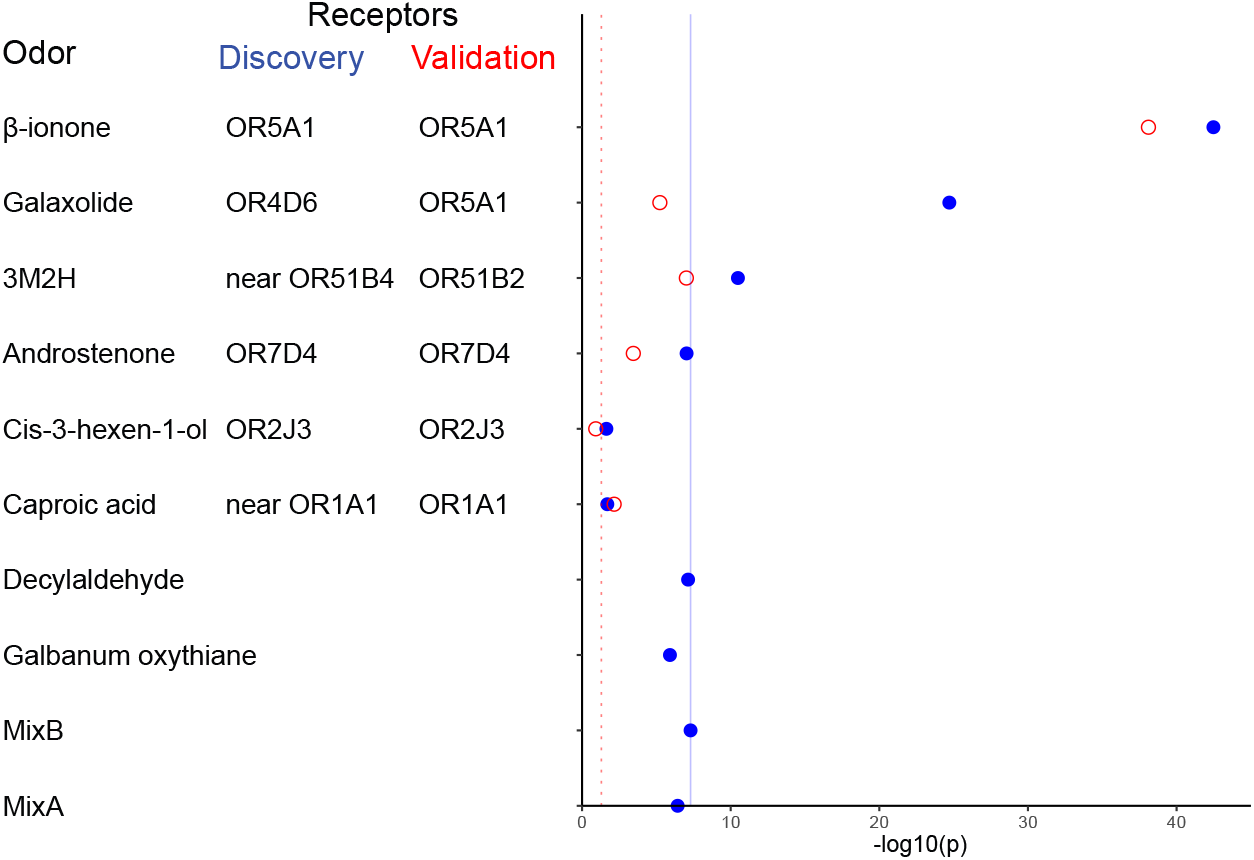
Top Associations Between Genetic Variation and Odor Perception. Each row represents the top SNPs associated with each odor in the discovery cohort (blue filled circles) and the replication cohort (red open circles). Listed next to each odor is the nearest gene to the top SNP for each cohort. There were two novel associations that reached genome-wide significance in the discovery cohort (p < 5×10^−8^, solid blue line): Galaxolide/rs1453541(M263T) and rs1453542 (S151T) (p<2.1×10^−25^, p<2.6×10^−25^) and 3M2H/rs3898917 (p<1.2×10^−11^). The discovery study replicated the associations from the literature (p<0.05, dotted red line) for *β*-ionone/rs6591536 (D183N) (p<3.3×10^−43^), androstenone/rs61729907 (R88W) and rs5020278 (T133M) (in both cases, p<1.1×10^−7^), and cis-3-hexen-1-ol/rs28757581 (T113A) (p<0.02), but not did examine the caproic acid/OR1A1 association. Other than *β*-ionone, no replication odors had associations that reached genome-wide significance. For these replication odors, the association shown is the top association from the LD-band surrounding the previously identified SNP. In the validation cohort (open red circles), we tested associations for the significant SNPs and surrounding LD-bands (±200kb) from the discovery study and previous literature. There were four significant associations in the validation study (p<0.05; red dotted line): *β*-ionone/rs6591536 (D183N) (p < 7.8×10^−39^), Galaxolide/rs591536 and rs7941591 (p<5.8×10^−6^), 3M2H/rs10837814 (L143F) (p<9.6×10^−8^), and androstenone/rs61732668(P79L) (p<3.5×10^−4^). The association for caproic acid/rs17762735 (p<6.9×10^−3^) is significant, but in the opposite direction predicted by the previous study.

### OR4D6 variant alleles M263T and S151T are associated with a decrease in Galaxolide intensity

In the discovery study, Galaxolide intensity perception was associated with an OR locus in chromosome band 11q12.1 (Fig. 2a-b). The two peak variants in open reading frames were both missense single nucleotide polymorphisms (SNPs) in OR4D6 (Fig. 2c): M263T (rs1453541, p<2.07×10^−25^) and S151T (rs1453542 p<2.59×10^−25^). The validation study confirmed that both OR4D6 SNPs correlated with the intensity of the higher of the two tested concentrations of Galaxolide (M263T p<9.08×10^−6^, S151T p<1.02×10^−5^, Fig. 2d). The two SNPs are in high linkage disequilibrium (LD): the variant allele of S151T is always co-inherited with the variant allele of M263T (Supplementary Table 2).

**Fig. 2.**
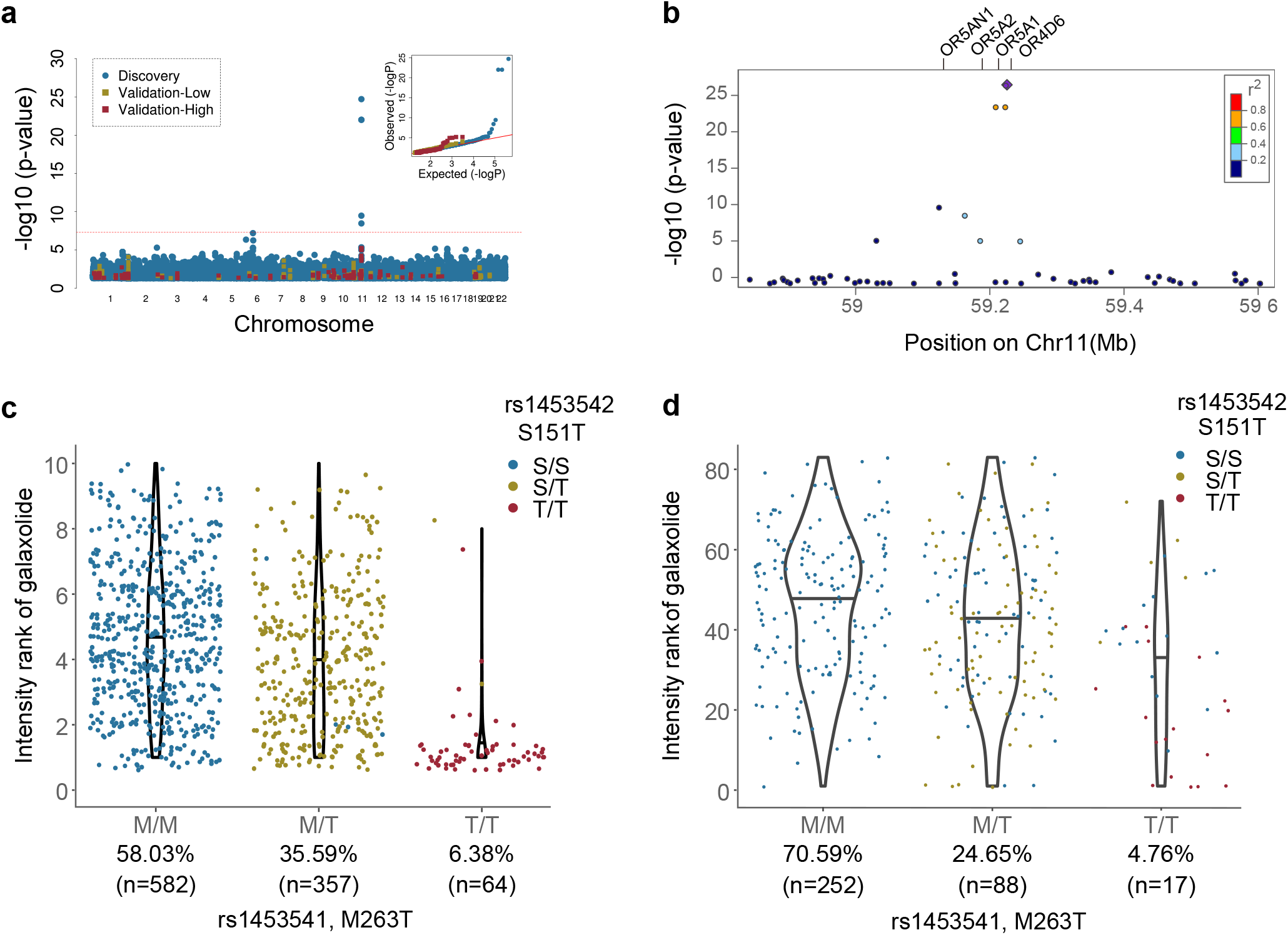
Galaxolide perception is associated with variation in chromosome band 11q12.1 in both cohorts, as shown by a) a Manhattan Plot of associations with the discovery study in blue and validation study in red (high concentration) and yellow (low concentration). The red line indicates the threshold for genome-wide significance (p<5×10^−8^). Inset: QQ plots from the discovery (blue) and validation ([high]=red, [low]=yellow) cohorts (Genomic Lambda: discovery=0.02; validation=0.90) show appropriate control for inflation due to population structure. b) The regional plot of discovery study associations indicates both the significance level and the recombination rate at the OR4D6 LD-band. Genetic variation in OR4D6 affects the perceived intensity of Galaxolide in c) the discovery cohort and d) the validation cohort (high concentration). The x-axis is ordered left-to-right with increasing number of variant alleles for the M263T variant, with population frequency of M263T indicated below the genotype. The points colored by S151T genotype suggest that in the validation cohort, S151T is driving the Galaxolide anosmia phenotype exhibited by those homozygous for the variant (T/T).

We examined the associations between Galaxolide and SNPs in other reported musk-activated ORs, including OR5AN1 (activated by muscone), OR5A2 (activated by all musk compound families), and OR1A1 (activated by nitro musks)(17, 18, 29). Of these ORs, only SNPs in OR5AN1 and OR5A2 were significantly associated with Galaxolide (p=2.98×10^−8^ and p=4.19×10^−17^, respectively; Supplementary Table 3). Since both of these SNPs are in strong LD with the novel signal discovered in OR4D6, we performed further analysis controlling for the top associated SNP in OR4D6, which did not reveal any signal reaching genome-wide significance. Additionally, only OR4D6 is in the credible set of the fine mapping analysis (Supplementary Table 4).

The leading role of OR4D6 is also supported by the meta-analysis of both cohorts where the two OR4D6 SNPs were the top two associations with Galaxolide intensity (M263T p<3.25×10^−29^, S151T p<4.63×10^−29^; Supplementary Data 2). The meta-analysis revealed no significant associations with Galaxolide pleasantness.

Based on the evidence from the above analyses we examined the effect size of the two SNPs in OR4D6 on Galaxolide intensity ratings. The S151T variant explains more of the phenotypic variance in Galaxolide intensity rankings (13.26% and 7.54% in discovery and validation cohorts, respectively) than M263T (12.84% and 4.74%). Variant homozygotes (T/T) ranked intensity lower than reference homozygotes (S/S or M/M) by an average of 33.3% and 17.1% (percentage of full scale) for M263T and 34.5% and 31.4% for S151T, for the discovery and validation cohorts, respectively (Fig. 2c-d).

To search for a mechanistic explanation for the observed associations, we tested high frequency (>5% frequency in validation cohort) OR haplotypes in high LD in this locus (OR4D6, OR5A1, OR5AN1, OR5A2; Fig. 2b; Supplementary Table 5). None of the ORs in the associated LD-block, or a consensus version of OR4D6 across 10 closely related species(10, 30) responded to Galaxolide in our assay.

### OR51B2 variant allele L134F is associated with increased 3M2H intensity

In the discovery study, 3M2H intensity perception was associated with an OR cluster in chromosome band 11p15.4 (Fig. 3a-b). The peak variant (rs3898917, p<1.20×10^−11^) is in a non-coding region in the LD band including OR51B2, and is in an expression quantitative trait locus (eQTL) affecting OR52A1(31). The validation study confirmed that this eQTL is correlated with 3M2H intensity ([low] p<7.89×10^−5^). There are a number of other associated variants in this LD band, but the only variant in the credible set of the fine mapping analysis (Supplementary Table 4) was a nonsynonymous missense SNP, rs10837814 (L134F) in OR51B2 (discovery p<6.57×10^−10^, validation p<9.60×10^−8^; Fig. 3c-d). The meta-analysis confirmed this as the only significant association with 3M2H intensity ([low] p<7.40×10^−16^), and found no further signal for pleasantness (Supplementary Data 2).

**Fig. 3.**
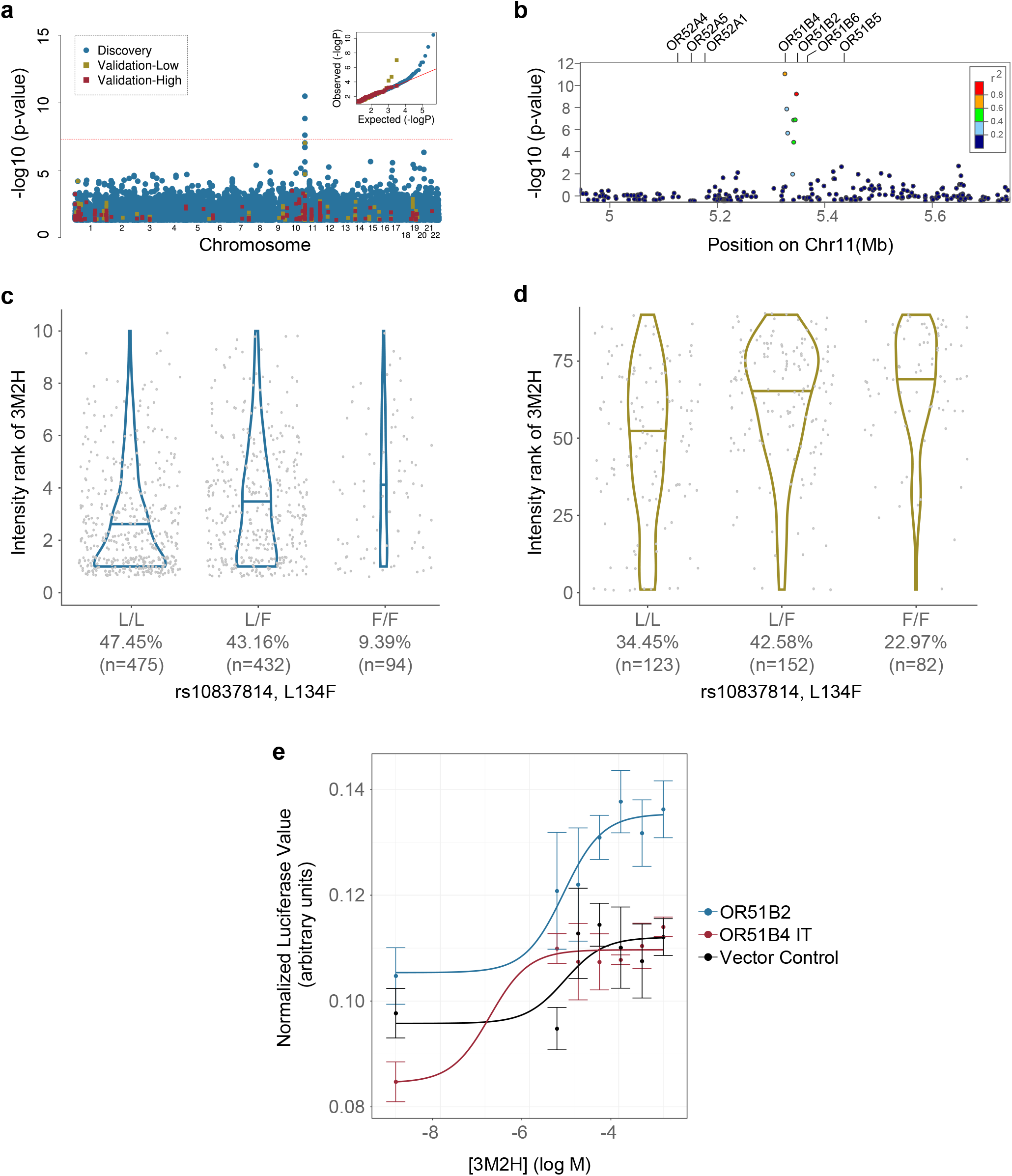
3M2H perception is associated with variation in chromosome band 11p15.4 in both cohorts as shown by a) a Manhattan Plot of associations with the discovery study in blue and validation study in red (high concentration) and yellow (low concentration). The red line indicates the threshold for genome-wide significance (p<5×10^−8^). Inset: QQ plots from the discovery (blue) and validation ([high]=red, [low]=yellow) cohorts (Genomic Lambda: discovery study=1.01; validation study=0.90) show appropriate control for inflation due to population structure. b) Regional plot of discovery study associations indicating both the significance level and the recombination rate at the OR51B2/4 LD-band. The variant L134F (rs10837814; OR51B2) was associated with the perceived intensity of 3M2H in the c) discovery and d) validation (low concentration) cohorts. The x-axis is ordered left-to-right for increasing number of variant alleles, with population frequency indicated below the genotype. e) The OR51B2 reference haplotype responds to 3M2H in a cell-based assay. The empty vector control (Rho) does not respond to 3M2H, nor do other receptors in the same LD-band such as OR51B4 (shown here; *DR model does not converge).

Given that the evidence from the meta-analysis and fine mapping analysis pointed to OR51B2, we examined the effect size of the L134F on 3M2H intensity ranking. The novel variant explains 4.13% and 9.97% of phenotypic variance in 3M2H intensity rankings in the discovery and validation cohorts, respectively. Variant homozygotes (F/F) ranked intensity higher than reference homozygotes (L/L) by an average of 12.8% in the discovery and 20.8% in the validation (Fig. 3c-d).

In order to further search for a mechanistic explanation for the observed associations, we used a cell-based assay to measure the response of high frequency (>5% frequency in the validation cohort) OR haplotypes in the associated locus (OR51B2, OR51B4, OR51B5, OR51B6; Fig. 3b; Supplementary Table 6), as well as in the eQTL-target locus (OR52A1, OR52A4, OR52A5; Supplementary Table 6), to 3M2H. OR51B2 responded to 3M2H (p<2.19×10^−5^; Fig. 3e). No other receptors in the OR51B2-associated locus or the eQTL-target locus responded to 3M2H (Supplementary Fig. 2).

### Replication of previously reported odor phenotype/OR associations

#### *β*-ionone/OR5A1

We replicated the association between *β*-ionone intensity perception and the missense SNP rs6591536 (D183N) in OR5A1 in the discovery cohort (p<1.84×10^−41^), the validation cohort ([high] p<7.80×10^−39^; Fig. 4a-b), and the meta-analysis (p<1.68×10^−74^). The pleasantness rank of *β*-ionone was also associated with D183N in the validation cohort ([high] p<5.53×10^−19^) and the meta-analysis (p<2.25×10^−9^), but not the discovery cohort (p=0.06). D183N was the top association with *β*-ionone, and other significant hits were all within the surrounding LD band (Supplementary Data 2). The variant D183N explains 21.6% and 31.9% of phenotypic variance in *β*-ionone intensity rankings in the discovery and validation cohorts, respectively. Variant homozygotes (N/N) ranked *β*-ionone intensity lower than reference homozygotes (D/D) by 20% and 38.3% in the discovery and validation cohort, respectively (Fig. 4c-d).

**Fig. 4.**
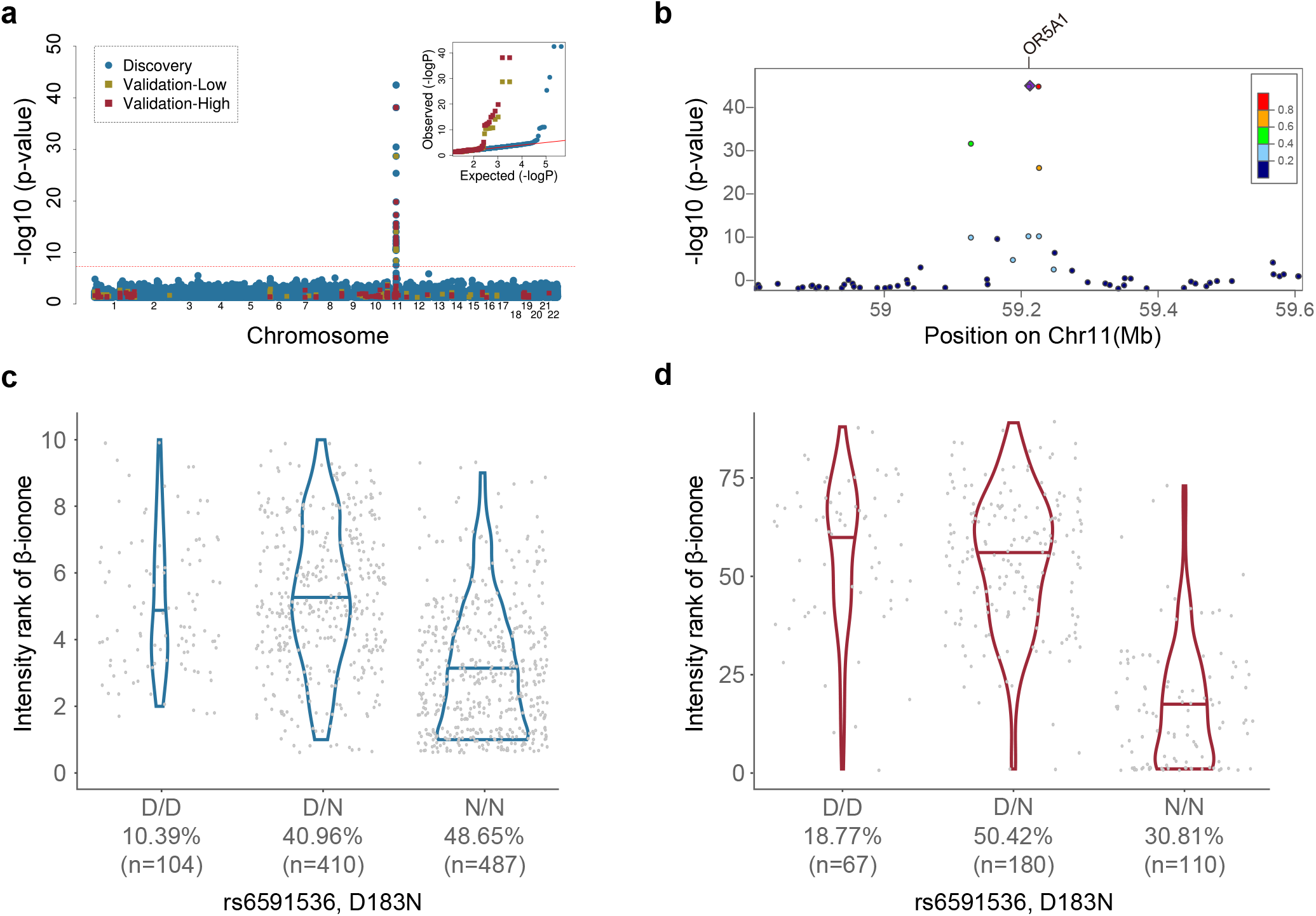
*β*-ionone perception is associated with variation in rs6591536 (OR5A1) in both cohorts, as shown by a) a Manhattan Plot of associations with the discovery study in blue and validation study in red (high concentration) and yellow (low concentration). The red line indicates the threshold for genome-wide significance (p<5×10^−8^). Inset: QQ plots from the discovery (blue) and validation ([high]=red, [low]=yellow) cohorts (Genomic Lambda: discovery study=1.00; validation study=0.94) show appropriate control for inflation due to population structure. b) Regional plot of discovery study associations indicating both the significance level and the recombination rate at the OR5A1 LD-band. The previously discovered D183N variant (rs6591536; OR5A1) also changes perceived intensity of *β*-ionone in our populations: c) discovery cohort, d) validation cohort (high concentration). The x-axis is ordered left-to-right for increasing number of variant alleles, with population frequency indicated below the genotype.

#### Androstenone/OR7D4

Androstenone perception has been previously associated with the RT/WM haplotype of OR7D4, which consists of two perfectly linked SNPs (rs61729907 (R88W) and rs5020278 (T133M)). This association was directly replicated in the discovery cohort for intensity (p< 1.07×10^−7^) and pleasantness (p< 9.56×10^−6^) ranking of androstenone. The validation cohort replicated the association for pleasantness (p<0.017), but not intensity (p=0.10). The meta-analysis found associations with RT/WM and both androstenone phenotypes (intensity p<6.25×10^−8^; pleasantness p<5.65×10^−7^). The validation cohort also confirmed the published effect of another OR7D4 SNP, rs61732668 (P79L), on androstenone perception (intensity p<3.53×10^−4^; pleasantness p<6.88×10^−3^). This SNP was not sequenced or successfully imputed in the discovery cohort and therefore could not be examined in the meta-analysis. In the discovery cohort, there was one novel association that reached genome-wide significance: rs117391865, an intronic SNP nearest the gene SYNE1 (p<1.35×10^−8^).

The RT/WM variants in OR7D4 predict 2.5% of phenotypic variance in androstenone intensity rankings in the discovery cohort. WM homozygotes ranked intensity lower than RT homozygotes by an average 20.6% (Supplementary Fig. 3). The P79L variant in OR7D4 predicts 2.8% of variance in androstenone intensity in the validation cohort. L homozygotes ranked intensity lower than P homozygotes by an average 25.6%.

These findings support previous literature: OR7D4 genetic variation has a consistent effect on androstenone perception, but explains only a small portion of the variance.

#### Cis-3-hexen-1-ol/OR2J3

The association between cis-3-hexen-1-ol intensity perception and rs28757581 (T113A) in OR2J3 was nominally replicated in the discovery cohort (p<0.02), but not in the validation cohort. The meta-analysis results suggest that this is an association at the low (p<0.03), but not high (p<0.08) concentration. In the meta-analysis, there are a number of associations in the LD band surrounding OR2J3 (including OR2W1 and OR2J1) at the p<0.05 significance level for pleasantness of cis-3-hexen-1-ol, and the top signal is rs3129158 in OR2J2 (p<2.3×10^−4^). No associations for cis-3-hexen-1-ol phenotypes reached genome-wide significance. Previous studies have not consistently demonstrated an association between cis-3-hexen-1-ol and OR2J3. Here we present evidence that some OR in this LD band may be involved in perception of cis-3-hexen-1-ol.

#### Caproic acid/OR1A1

We could not examine the previously published association between OR1A1 and caproic acid in the discovery study as this region was not sequenced or successfully imputed. Although rs17762735 in OR1A1 was associated with intensity in the validation study, the effect of the variant on the phenotype was in the opposite direction from the literature(10). There were no associations for pleasantness perception of caproic acid with OR1A1. No associations for caproic acid phenotypes reached genome-wide significance.

### There were no associations with aldehyde intensity or pleasantness

No significant signals were discovered for monomolecular aldehydes (decyl aldehyde and galbanum oxathiane) or either fragrances, MixA or MixB. The validation study did not examine these compounds.

### Derived alleles are ancient and tend to reduce odor intensity

Including the two novel SNPs reported in this study, we examined 29 SNPs that have been associated with odor perception in the literature (Table 1). In 24 of the 29 SNPs, the age of the derived allele was more than a hundred thousand years old (112,075-1,491,850 years), predating the population divergence times between East Asians and Europeans (∼55,000 years ago for East Asians and ∼41,000 years ago for Europeans)(32). Several SNPs existed in archaic humans and other non-human primates (Table 1). Based on the CMS score, there was no sign of natural selection for any of the 29 SNPs (Supplementary Fig. 4).

**Table 1.**
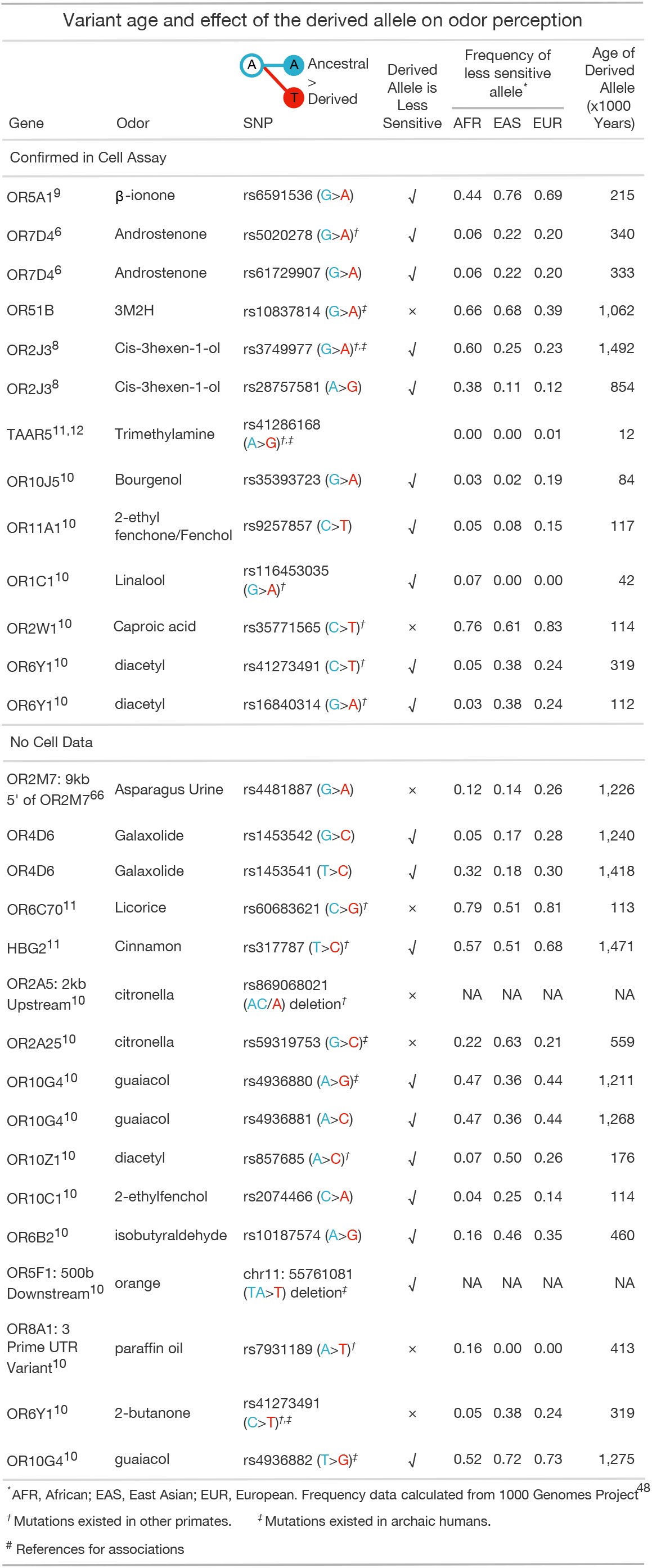
In a literature review, derived alleles corresponded with decreased sensitivity to odor in 21 out of 29 cases, and 11 out of 13 cases with functional validation. All but two variants predate the estimated ages of the East Asian and European population divergences.

| Variant age and effect of the derived allele on odor perception |  |  |  |  |  |  |  |
| --- | --- | --- | --- | --- | --- | --- | --- |
| Gene | Odor |  | Derived Allele is Less Sensitive | Frequency of less sensitive allele* |  |  | Age of Derived Allele (x1000 Years) |
|  |  | SNP |  | AFR | EAS | EUR |  |
| Confirmed in Cell Assay |  |  |  |  |  |  |  |
| OR5A1<sup>9</sup> | β-ionone | rs6591536 (G>A) | ✓ | 0.44 | 0.76 | 0.69 | 215 |
| OR7D4<sup>6</sup> | Androstenone | rs5020278 (G>A)<sup>†</sup> | ✓ | 0.06 | 0.22 | 0.20 | 340 |
| OR7D4<sup>6</sup> | Androstenone | rs61729907 (G>A) | ✓ | 0.06 | 0.22 | 0.20 | 333 |
| OR51B | 3M2H | rs10837814 (G>A)<sup>‡</sup> | × | 0.66 | 0.68 | 0.39 | 1,062 |
| OR2J3<sup>8</sup> | Cis-3hexen-1-ol | rs3749977 (G>A)<sup>†,‡</sup> | ✓ | 0.60 | 0.25 | 0.23 | 1,492 |
| OR2J3<sup>8</sup> | Cis-3hexen-1-ol | rs28757581 (A>G) | ✓ | 0.38 | 0.11 | 0.12 | 854 |
| TAAR5<sup>11,12</sup> | Trimethylamine | rs41286168 (A>G)<sup>†,‡</sup> |  | 0.00 | 0.00 | 0.01 | 12 |
| OR10J5<sup>10</sup> | Bourgenol | rs35393723 (G>A) | ✓ | 0.03 | 0.02 | 0.19 | 84 |
| OR11A1<sup>10</sup> | 2-ethyl fenchone/Fenchol | rs9257857 (C>T) | ✓ | 0.05 | 0.08 | 0.15 | 117 |
| OR1C1<sup>10</sup> | Linalool | rs116453035 (G>A)<sup>†</sup> | ✓ | 0.07 | 0.00 | 0.00 | 42 |
| OR2W1<sup>10</sup> | Caproic acid | rs35771565 (C>T)<sup>†</sup> | × | 0.76 | 0.61 | 0.83 | 114 |
| OR6Y1<sup>10</sup> | diacetyl | rs41273491 (C>T)<sup>†</sup> | ✓ | 0.05 | 0.38 | 0.24 | 319 |
| OR6Y1<sup>10</sup> | diacetyl | rs16840314 (G>A)<sup>†</sup> | ✓ | 0.03 | 0.38 | 0.24 | 112 |
| No Cell Data |  |  |  |  |  |  |  |
| OR2M7: 9kb 5' of OR2M7<sup>66</sup> | Asparagus Urine | rs4481887 (G>A) | × | 0.12 | 0.14 | 0.26 | 1,226 |
| OR4D6 | Galaxolide | rs1453542 (G>C) | ✓ | 0.05 | 0.17 | 0.28 | 1,240 |
| OR4D6 | Galaxolide | rs1453541 (T>C) | ✓ | 0.32 | 0.18 | 0.30 | 1,418 |
| OR6C70<sup>11</sup> | Licorice | rs60683621 (C>G)<sup>†</sup> | × | 0.79 | 0.51 | 0.81 | 113 |
| HBG2<sup>11</sup> | Cinnamon | rs317787 (T>C)<sup>†</sup> | ✓ | 0.57 | 0.51 | 0.68 | 1,471 |
| OR2A5: 2kb Upstream<sup>10</sup> | citronella | rs869068021 (AC/A) deletion<sup>†</sup> | × | NA | NA | NA | NA |
| OR2A25<sup>10</sup> | citronella | rs59319753 (G>C)<sup>‡</sup> | × | 0.22 | 0.63 | 0.21 | 559 |
| OR10G4<sup>10</sup> | guaiacol | rs4936880 (A>G)<sup>‡</sup> | ✓ | 0.47 | 0.36 | 0.44 | 1,211 |
| OR10G4<sup>10</sup> | guaiacol | rs4936881 (A>C) | ✓ | 0.47 | 0.36 | 0.44 | 1,268 |
| OR10Z1<sup>10</sup> | diacetyl | rs857685 (A>C)<sup>†</sup> | ✓ | 0.07 | 0.50 | 0.26 | 176 |
| OR10C1<sup>10</sup> | 2-ethylfenchol | rs2074466 (C>A) | ✓ | 0.04 | 0.25 | 0.14 | 114 |
| OR6B2<sup>10</sup> | isobutylaldehyde | rs10187574 (A>G) | ✓ | 0.16 | 0.46 | 0.35 | 460 |
| OR5F1: 500b Downstream<sup>10</sup> | orange | chr11: 55761081 (TA>T) deletion<sup>‡</sup> | ✓ | NA | NA | NA | NA |
| OR8A1: 3 Prime UTR Variant<sup>10</sup> | paraffin oil | rs7931189 (A>T)<sup>†</sup> | × | 0.16 | 0.00 | 0.00 | 413 |
| OR6Y1<sup>10</sup> | 2-butanone | rs41273491 (C>T)<sup>†,‡</sup> | × | 0.05 | 0.38 | 0.24 | 319 |
| OR10G4<sup>10</sup> | guaiacol | rs4936882 (T>G)<sup>‡</sup> | ✓ | 0.52 | 0.72 | 0.73 | 1,275 |
| * AFR, African; EAS, East Asian; EUR, European. Frequency data calculated from 1000 Genomes Project.<br>† Mutations existed in other primates. ‡ Mutations existed in archaic humans.<br># References for associations |  |  |  |  |  |  |  |

\* AFR, African; EAS, East Asian; EUR, European. Frequency data calculated from 1000 Genomes Project<sup>48</sup>
<sup>†</sup> Mutations existed in other primates. <sup>‡</sup> Mutations existed in archaic humans.

In 21 out of 29 examined cases the derived allele was less sensitive to odors (72.4%; p<0.01). 13 of these 29 SNPs have been functionally validated by cell assay. Of these 13, there were 11 cases where the derived allele associated with decreased odor sensitivity (84.6%; p<0.01).

## Discussion

We conducted a genome-wide association study using ten odors and found novel associations for OR4D6 with the musk odor Galaxolide, and OR51B2 with 3M2H. In addition, we replicated previous associations between OR5A1/*β*-ionone, OR7D4/androstenone, and OR2J3/cis-3-hexen-1-ol. Furthermore, we have shown that these genotype/phenotype associations are stable across populations and robust to differences in methods, including odor concentration and delivery method. Previous genotype/phenotype studies have tended to focus on variation in olfactory receptors, however differences in olfactory perception could be driven by genetic variation in other proteins involved in odor signal transduction, such as olfactory axon guidance molecules, odor-modifying enzymes, or odor transport proteins. Despite our genome-wide search, the peak associations were largely located within olfactory receptor loci, suggesting that differences in olfactory perception caused by genetic factors are frequently driven by changes in the receptors.

### OR4D6 variation drives differences in perception of Galaxolide, but multiple receptors are involved in musk perception

Musks are a chemically diverse set of compounds that are defined by their common perceptual quality; however variation in perception of intensity of different musk odors across individuals(15, 16) suggests several receptors or groups of receptors may have a contributing role. The musk family, therefore, provides us with an opportunity to study the convergence of perceptual features of odors through differential receptor activation in the olfactory code. Prior to our study, there were four human olfactory receptors that responded to musks in cell culture(17, 18, 29), but their influence on olfactory perception is unknown. Here we identified a fifth musk receptor, OR4D6, where genetic variation associated with differences in perception of the polycyclic musk, Galaxolide. This is the first behavioral evidence that any human olfactory receptor plays a role in musk perception.

Other receptors that may be involved in musk perception have shown specificity for a particular musk or musk chemical family. Mice with a genetically deleted Olfr1440 (MOR215-1) were unable to find muscone in an odor-finding task(18), suggesting that the receptor is necessary for detection of the polycyclic musk muscone. The human ortholog of Olfr1440, OR5AN1, has relatively high affinity for several macrocyclic and nitro musk compounds in a heterologous cell-based assay(17, 18). Screening with this cell-based assay uncovered two other putative human musk receptors, OR1A1 and OR2J3(18), which respond only to nitro musks, but not Galaxolide or other polycyclic musks. There is also recent evidence for a broadly tuned musk receptor, OR5A2, which is activated by musks from all four tested structural classes in vitro(29). Together, the existence of musk-specific or musk-family specific, as well as broadly tuned musk receptors suggests that musks activate separate, but potentially overlapping, sets of receptors.

Here, we have identified the first case where genetic variation in a receptor is associated with musk perception in humans. Although OR4D6 is the top association with the Galaxolide intensity phenotype, it is in high linkage disequilibrium with two previously identified musk receptors: OR5AN1, and OR5A2. Due to solubility issues with Galaxolide in our cell-based assay, we were unable to provide functional evidence for OR4D6; however, several pieces of evidence support the idea that genetic changes in OR4D6 are driving the phenotypic difference in Galaxolide intensity: *1*. In both cohorts OR4D6 is a stronger predictor of Galaxolide intensity than OR5A2 and OR5AN1, which are not significantly associated with the phenotype after controlling for the top variant, and *2*. With few exceptions, participants homozygous for the OR4D6 variant are unable to smell Galaxolide.

OR4D6 is a strong candidate for the mechanism underlying specific anosmia to Galaxolide, suggesting that it is possible for a single receptor to represent the musk percept. We do not know if OR4D6 contributes to perception of other musk compounds, but given the in vitro evidence for other musk receptors and behavioral data that suggest those with Galaxolide anosmia are still able to smell other musk compounds(15, 16), it is unlikely to be solely responsible for the perception of the musk quality percept. OR4D6, OR5AN1, and OR5A2 are prime targets for future work on musks, which can lead more broadly to understanding how activation of different combinations of receptors results in highly similar percepts.

### OR51B2 variation drives differences in the perception of human body odor component 3M2H

Trans-3-methyl-2-hexenoic acid (3M2H) has been described as the ‘impact odor’ for body odor arising from the underarms, meaning that it is a highly abundant volatile compound and its quality as a monomolecular odorant is the same as the characteristic quality of body odor(23). Specific anosmia to 3 M2H has been reported in several studies with rates ranging from 21-25% of the population(23, 25, 26). Based on its key role in body odor character, it is likely that anosmia to 3M2H alters body odor perception, although it does not eliminate the ability to smell body odor, as there are other reported volatile compounds present in underarm odor(23, 26, 33).

Here we found that OR51B2 was associated with 3M2H intensity, and responded to 3M2H in a functional assay, suggesting that OR51B2 drives differences in perception of 3M2H intensity. Through this mechanism, we therefore predict that OR51B2 genotype will impact the perception of body odor. OR51B2 could be a target for future studies interested in malodor blocking, or discovering the mechanisms underlying social communication from body odor.

### Associations between OR5A1/*β*-ionone, OR7D4/androstenone, and OR2J3/cis-3-hexen-1-ol are replicated in an East Asian population

Here for the first time in an East Asian population, we replicated previous phenotype/genotype associations between OR5A1/*β*-ionone/, OR7D4/androstenone, and OR2J3/cis-3-hexen-1-ol, but failed to replicate the OR1A1/caproic acid association. The N183D OR5A1 variant has now been associated with decreased intensity perception of *β*-ionone in several studies(9, 34), as well as verified in a cell-based assay(9). In both the discovery and the validation cohorts, the *β*-ionone intensity phenotype had the highest overall effect size, showing this association is not only robust to differences in methods, but has also been replicated across multiple populations.

In the discovery cohort, we replicate the association between RT/WM haplotypes of OR7D4 and androstenone perception(6), that has been replicated in two other populations(35, 36). Although the validation study did not replicate the association with androstenone intensity and RT/WM, it did replicate the association with androstenone pleasantness, as well as the previously discovered association between P79L in OR7D4 and androstenone perception(6). The lack of signal for intensity perception and RT/WM in the validation study could be due to differences in odor delivery method or concentration of the odor. Overall, the evidence here continues to support the role of OR7D4 in androstenone perception.

Here we nominally replicated the association between cis-3-hexen-1-ol and OR2J3(8). The smaller signal here is not surprising, as this association has failed to replicate in two other studies(10, 34). The original discovery of this association measured the detection threshold of cis-3-hexen-1-ol, while the two studies that failed to replicate the original association measured intensity rankings. Since the set of receptors that associate with variation in olfactory perception differs across concentrations(10), this could explain why the cis-3-hexen-1-ol/OR2J3 association failed to replicate previously and only has a small signal here.

A previous study found an association between OR1A1 and caproic acid(10). This failed to replicate in the validation study, and had no direct replication provided from the discovery study. The discovery study did have data on variants in the locus surrounding OR1A1 that had an association signal with caproic acid p<0.05. The role of OR1A1, or perhaps another OR in this region, in the perception of caproic acid is still unclear.

Although this is the first study to examine a large East Asian population (as opposed to a majority-European population), the majority of associations replicate across different populations. Summarizing the variants that alter odor perception, including those reported in past literatures, our analysis of evolutionary age found 25 of 27 variants predated the population divergence between East Asians and Europeans (∼55,000 years ago for East Asians and ∼41,000 years ago for Europeans)31. These variants are generally present in both populations at relatively high frequencies. On the other hand, the two more recently derived variants (S95P in TAAR5 and A67V in OR1C1) have very low minor allele frequencies in both East Asian and European populations, suggesting population specific variants that alter odor perception are rare.

### Variation in the perception of aldehydes does not associate with olfactory receptors

A study in a large non-homogenous population from New York, NY, tested perceptual differences of 15 fragrances between self-reported demographic groups. Of all the odors tested, the largest difference found was in aldehydes, and all three tested aldehydes (decyl aldehyde, nonyl aldehyde, and undecanal) were significantly different, such that the Asian population perceived aldehydes to be more intense than the Caucasian population(28). The follow-up study pursuing the genetic underpinnings of these differences did not identify any associated ORs(10), and even here, in a larger cohort with a genome-wide search, there were no associations.

The lack of genetic evidence here may also be due to the involvement of multiple ORs, reducing our power to detect specific associations; or the perceptual variation may be due to cultural, social, or other factors that are not genetic in nature. The more recent evolutionary age of population-specific variants may play a role, as this these types of odor analyses have discovered mostly ancient variants, with only two odor-associated SNPs that appeared after the East Asian and European population divergences. This suggests that increasing population size, regardless of diversity, may be necessary to discover more recently derived SNPs with lower minor allele frequency.

### Degeneration of olfactory receptor gene repertoires in primates

Compared to many non-primate mammalian species, primates have fewer intact olfactory receptor genes both in absolute number and by percentage(37). While previous analyses have been restricted to pseudogenes, recent analyses of the functional consequences of missense mutations allow for a more detailed examination. We found that in 72% of reported OR gene/olfactory phenotype associations reported in the literature (85% with functional validation), derived alleles predicted lower perceived intensity than ancestral alleles. While this study was not designed to directly address this hypothesis and may suffer from selection bias, these data support the hypothesis that the primate olfactory gene repertoire has degenerated over time. The functional implications of this degeneration remain unclear(38, 39).

### Large genetic databases can be used to understand OR function, a proxy for general protein function

In the discovery study, we have the benefit of measuring olfactory phenotypes in a cohort where genome-wide genotyping had already been conducted, giving us the statistical power of a large population without the time or expense. Given the increasing number of open databases of sequencing data, this method is becoming a more reasonable possibility for easily testing genotype/phenotype associations.

Olfaction is an excellent use of this new resource because of the ease of understanding the functional output of genetic variation in the protein. The human olfactory system has both robust assays to test the behavioral output of these proteins (psychophysics/rating odors)(5, 6, 10) and an established method for directly testing protein function in cells (heterologous cell-based assay)(40, 41). Genetic variation provides a strong tool for exploring olfactory coding and sheds light on how complex systems integrate information from variable sensors.

## Supporting Information (SI)

Supporting information is supplied in a separate.pdf

### SI Figures

Supporting Figures are included with Supporting Tables in a separate.pdf

**SI Fig.1** Phenotype Distribution

**SI Fig.2** 3M2H/OR51B1 LD-band Cell-based Assay Results

**SI Fig.3** Androstenone Intensity by OR7D4 RT/WM genotype in Discovery Cohort

**SI Fig.4** Natural Selection Results

**SI Fig.5** PCA of Population Structure for Discovery, Validation, and 1000 Genomes Data

### SI Tables

Supporting Figures are included with Supporting Tables in a separate.pdf

**SI Table 1** Phenotype Heritability

**SI Table 2** Frequency of Linked SNPs in OR4D6 Associated with Galaxolide Perception

**SI Table 3** Associations Between Galaxolide and SNPs of Other Reported Musk-Related ORs

**SI Table 4** Fine Mapping Analysis

**SI Table 5** OR Haplotypes Tested in the Cell-based Assay for Activation by Galaxolide (OR4D6 Cluster)

**SI Table 6** OR Haplotypes Tested in the Cell-based Assay for Activation by 3M2H (OR51B2 Cluster)

### SI Datasets

Supporting datasets are provided in separate files.

**SI Data 1.** Significant Discovery Cohort Associations (p< 5×10^−8^).

**SI Data 2.** Meta-Analysis Results.

## ACKNOWLEDGMENTS

This work was supported by the National Key Research and Development Project (Grant No. 2018YFC0910403), the National Natural Science Foundation of China (Grant No. 91631307), Shanghai Municipal Science and Technology Major Project (Grant No.2017SHZDZX01), CAS Youth Innovation Promotion Association (Grant No. 2020276), the National Institutes of Health (Grant R01 DC013339), and in part by the National Center for Advancing Translational Sciences Clinical and Translational Science Award program (grant UL1 TR000043). The discovery study was funded by Unilever R&D (the Netherlands). A portion of the work (validation study) was performed at the Monell Chemosensory Receptor Signaling Core, which was supported in part by the National Institute on Deafness and Other Communication Disorders (Core Grant P30 DC011735). We acknowledge the contributions from Young de Graaf from Unilever for making the odorant solutions in the discovery study; Marcia Knoop and the sensory panel from Unilever for isointensity testing; Tom Salmon from Unilever, for his help in selecting odorant stimuli and fragrances; and David Gunn from Unilever for helpful comments throughout the project.

## Author Contributions

SW and JM conceived the project, provided resources, and edited the manuscript. BL collected samples of the discovery cohort. BL and QP contributed to analysis of the discovery cohort data, meta and evolutionary. MK contributed to analysis of the validation cohort, functional validation experiments. FL, AK, MS contributed to the design of the project. MK prepared the initial draft of the manuscript with materials from BL and QP All authors discussed and made contributions to the final version.

## Materials and Methods

### Study cohorts and participants

The discovery cohort comprised 1003 participants between the ages of 18 and 55, from a Han Chinese population collected in Tangshan, China. The research was conducted under approval of the Ethics Committee of Chinese Academy of Sciences (Shanghai, China). The validation cohort comprised 364 participants between the ages 18-50, from a diverse population collected in New York, New York, USA. Psychophysics experiments were conducted under approval from the IRB at Rockefeller University (New York, NY).

Both cohorts excluded participants with medical conditions that affect the sense of smell, specifically: smoking, recreational drug use, brain surgery or head trauma that required hospitalization, chronic nasal issues (allergic, tumoral, infectious or inflammatory disease), history of endoscopic nasal or sinus surgery, any neurodegenerative disease, any upper respiratory infection that altered the sense of smell and/or taste for more than 1 month, cervicalgia or other neck diseases, history of radiation or chemotherapy, alcoholism, current sinus or upper respiratory infection, seasonal allergic rhinitis or acute rhinosinusitis, and use of medications that interfere with the sense of smell.

### Odor Delivery

Discovery cohort participants were tested using felt-tip pens (100.2 mm length, diameter 7.7 mm; ETRA, KönigsbachStein, Germany) containing an absorbent material loaded with 1 mL of liquid odor. Each pen was used for no more than 50 participants before being discarded. After preparation, individual sticks were used within 2 months.

Validation cohort participants smelled 20 ml amber glass vials filled with 1 mL of odor. The vials were presented in a double-blind manner, labeled only with barcodes, to prevent experimenter bias.

### Phenotyping

Discovery cohort participants smelled 11 odors (10 unique and one repeat) and verbally rated the intensity and pleasantness on a 100-point scale. For each odor, a unique set of 100 participants rated the stimulus twice so we could measure the test-retest reliability. Validation cohort participants also smelled each stimulus and rated intensity and pleasantness on a 100-point computerized sliding scale. The participants smelled 46 odors at one concentration, 26 odors at two concentrations, three odors at five concentrations, and three solvents for a total of 90 stimuli, ten of which are reported here (six odors: four at low and high concentration and two at high concentration only). Participants smelled stimuli in the same order to facilitate comparisons across participants, and every stimulus was presented twice.

In both cohorts, in order to normalize for scale usage across raters, intensity and pleasantness ratings for each participant were ranked from 1 to 10, or 1 to 90, such that the odorant with the lowest rated intensity was ranked at 1, and the odorant with the highest intensity was ranked 10 or 90, depending on the total number of stimuli (Supplementary Fig. 1)(6). The change in ranking metric was calculated as a percentage of the number of ranks changed over the total number of ranks in the scale (10 or 90), in order to directly compare changes between the cohorts.

To measure the within-subject reliability of ratings, we calculated the Pearson’s correlation between duplicate stimuli. In the validation cohort, 4 participants were removed due to poor performance (test-retest correlation <= 0). Individual performance could not be examined in the discovery cohort, as only one stimulus was repeated for each participant.

### Odor concentration and preparation

The discovery cohort included 8 monomolecular odorants (androstenone, *β*-ionone, caproic acid, cis-3-hexen-1-ol, Galaxolide, trans-3-methyl-2-hexenoic acid (3M2H), decyl aldehyde, and galbanum oxathiane) and 2 odor mixtures (MixA and MixB) all prepared by Unilever. We used isointense concentrations of the ten odors that were diluted in either propylene glycol or MCT (medium chain triglycerides) (Table 2). To determine the concentration, 14 expert panelists rated intensity of ten odorants at three different concentrations (except 3M2H and androstenone which were rated at 2 concentrations) that were pre-selected to cover a range from weak to strong. Panelists rated intensity on a scale from 0-15, using a range of concentrations of citric acid for reference. Ratings were significantly difference between all concentrations of odors, except for androstenone, for which we chose the higher concentration. For each odor, we chose the concentration that was closest to an intensity of 7, with the exception of two odors (caproic acid and MixA) for which an original concentration did not result in a rating near 7. For these odors, we extrapolated the concentration that would result in an intensity rating of 7 from the other intensity ratings.

**Table 2.** Concentrations of Odors from the Discovery and Validation Studies

| Odor | Discovery Cohort |  | Validation Cohort |  |  |
| --- | --- | --- | --- | --- | --- |
|  | Dilution | Solvent | Dilution (high/low) | Solvent | Alternate Name |
| β-ionone | 50X | Propylene Glycol | 1/10,000<br>1/400,000 | Paraffin Oil | 3E-4-(2,6,6-Trimethylcyclohex-1-en-1-yl)but-3-en-2-one |
| 3M2H | 0.1 g/mL | Propylene Glycol | 1/100<br>1/20,000 | Paraffin Oil | trans-3-methyl-2-hexenoic acid |
| Galaxolide | 2X (50% in diethylphthalate) | MCT (medium chain triglycerides) | 1/10<br>1/1000 (weight/volume) | Paraffin Oil | 1,3,4,6,7,8-hexahydro-4,6,6,7,8,8-hexamethylcyclopenta[g]-2-benzopyran |
| Cis-3-hexen-1-ol | 200X | Propylene Glycol | 1/100,000<br>1/250,000 | Paraffin Oil | NA |
| Decylaldehyde | 1000X | Propylene Glycol | NA | NA | decanal |
| Androstenone | 1.42 mg/mL | Propylene Glycol | 1/1000 weight/Volume | Propylene Glycol | 5α-androst-16-en-one |
| Caproic acid | 500X | Propylene Glycol | 1/1,000,000 | Paraffin Oil | hexanoic acid |
| Galbanum oxathiane | 5000X | Propylene Glycol | NA | NA | (2R,4S)-2-methyl-4-propyl-1,3-oxathiane |
| MixB | 1000X | Propylene Glycol | NA | NA | NA |
| MixA | 400X | Propylene Glycol | NA | NA | NA |

The validation cohort includes data from the following six odors: androstenone, *β*-ionone, cis-3-hexen-1-ol, caproic acid, Galaxolide, and 3M2H. The aldehydes and fragrances were not measured in the validation study. High and low concentrations of odors were intensity-matched to 1/1,000 and 1/10,000 dilutions of 1-butanol, as determined by rankings from a panel of 13 individuals. Odors were presented at both concentrations, except for androstenone and caproic acid, which were given at concentrations based on previous studies(6, 10). Odors were diluted in paraffin oil or propylene glycol (Table 2).

Due to different delivery methods in each cohort, the concentrations of these six compounds cannot be directly compared to the concentrations in the discovery study(42, 43).

### Genotyping

Discovery Cohort: Genomic DNA was extracted from blood samples using the MagPure Blood DNA KF Kit. All samples were genotyped using the Illumina Infinium Global Screening Array that analyzes over 710,000 SNPs. It is a fully custom array designed by WeGene (https://www.wegene.com/).

Validation Cohort: Genomic DNA (gDNA) was extracted from saliva samples using the Oragene Discover 2mL kit and protocol. Library prep (using Agilent SureSelect XT2 kit) and targeted sequencing were performed by CAG sequencing core (Children’s Hospital of Philadelphia Research Institute, Philadelphia PA). Custom Agilent SureSelect targets were designed (eLID# 3028991) for 418 ORs and 290 olfactory-related genes, including other odorant receptors (i.e. TAARs, MS4A) and related enzymes (i.e. CYP). The Illumina HiSeq platform was used to perform paired-end sequencing with a read length of 2×125 basepairs on 364 participants.

### Variant calling and quality filtering

Discovery Cohort: Sequences were aligned to genome build GRCh37/hg19 and genotypes were called using Genome Studio v2.0(44). To control for genotype quality, we implemented exclusion criteria using PLINK v1.90b6.9(45). No people were removed due to >5% missing data or failure of X-chromosome gender concordance check. We excluded SNPs that had >2% missing data (14,385 variants removed), a minor allele frequency (MAF) <1% (251,918 variants removed), or a deviation from Hardy-Weinberg (HW) equilibrium (p<1×10^−5^)(46) (1,149 variants removed), leaving 433,485 SNPs from 1003 individuals for genome wide association analysis. SNP phasing was performed with Eagle v2.4(47) using 1000G Phase 3 V5 (GRCh37/hg19) EAS as the reference panel(48). We conducted imputation on the 433,485 phased SNPs using Minimac4, and obtained a total of 45,843,286 variants. We then re-ran genotype quality control steps and filtered out 54 variants missing >2% genotype data, 27,361 variants with a deviation from HW equilibrium (p<1×10^−5^)(46), and 37,772,956 variants due to MAF threshold (MAF>0.01), leaving 8,042,915 variants for association analysis.

Validation Cohort: Genotypes were called using a pipeline that follows recommended ‘best practices’ by the Broad Institute(49, 50), and as previously reported(10). Sequences were aligned to GRCh37/hg19 genome build using BWA(51), and alignment, genotype quality and variant calling steps were performed using Picard Tools(52, 53). SNP phasing was performed with SHAPEIT V2.r900(54), and OR haplotypes were assembled using a custom R script. Of the original 18,611 variants called, quality control measures filtered out 1,488 SNPs that were missing genotype data at a frequency >5% or deviated from Hardy-Weinberg equilibrium (p<1×10^−5^)(46). An additional 14,078 variants were removed due to minor allele frequency (MAF>0.05), leaving 3,045 variants for association analysis. Three individuals were excluded due to > 5% missing data, leaving 357 participants remaining for genotype/phenotype analysis. For one region of the genome (chromosome band 11p15.4) the discovery study found significant association in a non-coding region. This region was not sequenced in the validation study, which focused on open reading frame variants, so we imputed 147,613 SNPs in this region (11:79438 to 11:249222325, hg19).

### Population structure analysis

We combined the discovery and validation datasets in order to visualize and quantify differences in the two study populations. We performed principle component analysis (PCA) using 1,018 linkage disequilibrium-pruned (r2<0.2) SNPs from the combined discovery (n=1003), validation (n=357), and 1000 Genomes Project (phase 3, 271 participants: 97 CHB, 86 CEU, and 88 YRI) datasets(48). We calculated centroids for each population using the first two eigenvectors. The distances between populations were measured by Euclidian distance of the centroids. The distances within a population were measured by averaging the Euclidian distance between each point (participant) and the centroid in the population.

The first two principle components explained 52% and 23% of the genetic variance (Supplementary Fig. 5). Genetic distance analysis confirmed that the discovery population overlapped with the CHB (Han Chinese population from the 1000 Genomes Project(48)) (mean distance to CHB=0.001, CEU=0.07, YRI=0.07), whereas the validation study population was distributed between different superpopulations (mean distance to CHB=0.05, CEU=0.05, YRI=0.05) (Supplementary Fig. 5). The mean distance between any two participants within a study cohort is smaller in the discovery population (mean distance = 0.007) than in the validation population (mean distance = 0.047), confirming that the discovery population is more homogeneous than the validation population (p<2.2×10^−16^; Supplementary Fig. 5).

### Association analysis

Discovery Cohort: To control for population stratification, we identified the top 10 genetic eigenvectors to use as covariates by performing PCA on 143,988 LD-pruned (r2<0.2) SNPs from the 1003 participants of the discovery cohort using Plink v1.90b6.9(45, 55).

Using PLINK (v1.90b6.9)(45), we performed genome-wide association analyses of 8,042,915 SNPs against 20 ranked phenotypes (intensity and pleasantness of 10 odors) under an additive linear model including age, sex, and the top ten genetic eigenvectors as covariates. Associations were significant if they passed the genomewide significance threshold (p<5×10^−8^). For loci of interest, we calculated linkage disequilibrium using LocusZoom(56) using the genome build from hg19/1000 genomes Nov 2014 ANS. We estimated the heritability of each perception phenotype explained by LD-pruned SNP set (143,988 SNPs with r2<0.2) using GCTA software (v1.93.0 beta)(57, 58).

Validation Cohort: To determine the top 10 genetic eigenvectors(55, 59) for the validation study, we conducted PCA on 10,927 LD-pruned (r2<0.05) SNPs with <5% missing genotypes and in HW equilibrium (p<1e-5) (but without excluding for MAF) from 361 people (including participants later excluded for poor phenotype data) using the R/Bioconductor package SNPRelate(60).

We performed genetic association analysis using PLINK (v1.90b5)(45) to test additive linear models for the 3,045 SNPs from quality control steps and the 78,904 SNPs from the imputed region against each of the 20 phenotypes of interest (intensity and pleasantness of six odors at one or two concentration each; see Table 2) with the top ten genetic eigenvectors as covariates. For significant loci from the discovery study we set alpha=0.05. For these loci of interest, we calculated linkage disequilibrium using LocusZoom(56) with the genome build from hg19/1000 genomes Nov 2014 EUR.

Combined Cohorts: We conducted a meta-analysis of the discovery and validation cohorts with METAL(61), which combines weighted p-values, weighted by sample size, across studies while taking into account and direction of effect.

### Fine mapping analysis

We conducted the fine-mapping analysis by leveraging functional annotation data (GenCode.exon.hg19) and LD information in the discovery and replication cohorts(62). We assumed a single causal variant at each locus, examined the SNPs within 200kb upstream and downstream of the top variant, and calculated the posterior probabilities using PAINTOR to determine the 99% credible set.

### Olfactory receptor cloning and haplotypes

To determine functional consequences for the identified SNPs in the olfactory receptors (ORs) and nearby receptors in high linkage disequilibrium, we tested activation of specific haplotypes of the associated ORs, as well as nearby ORs in the same LD-band. We have a large library of variant and reference haplotypes of ORs that we can use for testing differential response of receptor variants in the cell-based Luciferase assay. To supplement our library, we ordered and subcloned an OR4D6 consensus sequence into the vector pCI-RHO (GenScript). pCI-Rho (Promega) contains the first 20 amino acids of human rhodopsin(63). Using a consensus version of a receptor can improve surface expression in a heterologous cell-based assay where the original receptor is not expressed(10, 30).

We created a consensus sequence for OR4D6 using orthologs found in Homo sapiens, Gorilla gorilla, Pan paniscus, Pan troglodytes, Pongo abelii, Macaca mulatta, Mandrillus leucophaeus, Callithrix jacchus, Microcebus murinus, Rattus norvegicus, and Mus musculus. We aligned the orthologs using the online version of MAFFT version 7(64), and determined the most common amino acid at each position for the open reading frame of OR4D6. The consensus amino acid sequence was printed by GenScript and subcloned into the pCI-Rho vector (Promega).

### Luciferase assay

We used a heterologous cell-based assay to determine the functional changes caused by different OR haplotypes for our two novel associations, as has been previously described(10, 40, 41).

*Transfection*: Using the Dual-Glo Luciferase Assay System (Promega). We transfected Hana3A cells with our OR of interest, firefly luciferase driven by a cyclic AMP response element (CRE) promoter, and Renilla luciferase driven by a constitutively active SV40 promoter, RTP1S63, and M3-R(65).

*Stimulation:* Approximately one day after transfection, we stimulated cells by adding the odor in a 3-fold dilution series in CD293. Each concentration was run in triplicate, including the empty vector negative control. Stock odors were kept at 1M in DMSO and diluted in CD293 to the highest applied concentration of 1mM. Four hours after adding odor to cells, we read the luminescence output using a Synergy 2 plate reader (BioTek). Luciferase values were normalized by Renilla luciferase to control for transfection efficiency and cell death, and then averaged across the triplicate readings.

*Analysis:* The normalized luciferase values were fit to a three-parameter sigmoidal curve with a fixed slope (slope=1). We considered a receptor to be activated by an odorant if the response passed three tests: 1) the standard error of the logEC50 was less than one log unit, 2) The 95% confidence intervals for the top and bottom parameters of the curve did not overlap, and 3) The dose response curve from the OR-transfected cells was significantly different from the negative control (empty vector), as calculated by the extra sum-of-squares test. Data analysis was performed using GraphPad Prism (Version 8).

### Evolutionary analysis

We accessed the dbSNP database (https://www.ncbi.nlm.nih.gov/snp/) to determine the derived and ancestral alleles for our two novel SNP associations and 29 SNPs with previously reported odor phenotype associations(5–11, 66, 67). To our knowledge, this included all previously published associations between a SNP and an olfactory phenotype, exclusive of haplotype associations where direction of effect from individual SNPs could not be determined. We estimated the age of derived alleles using a Genealogical Estimation of Variant Age (GEVA) model (https://human.genome.dating/)(68). We checked if these mutations existed in archaic humans (i.e. Neandertal and Denisova) or in other primates using publicly available sequences (https://genome.ucsc.edu/Neandertal/,http://cdna.eva.mpg.de/denisova/)(69) and UCSC database (https://genome.ucsc.edu/). We also tested whether these SNPs were under positive selection using the Composite of Multiple Signals (CMS) method(70). To examine the relationship between derived alleles and a decrease in odor intensity perception, we performed a one sided two-proportions z-test (R version 6.3.1).

## Data availability

The GWAS summary statistics were deposited in the National Omics Data Encyclopedia (http://www.biosino.org/node/), and are available upon reasonable request under the Project ID: OEP001806. Individual-level genotype and phenotype data are not publicly available owing to them containing information that could compromise research participant privacy or informed consent. All other data are contained in the article file and its supplementary information or available upon reasonable request to the corresponding authors.

