## Supplemental Figures and Tables for "From musk to body odor: decoding olfaction through genetic variation"

Supplemental Information: Preprint

Bingjie Li[1,2,3] Marissa L. Kamarck[1,4,5] Qianqian Peng[1,2] Fei-Ling Lim[6]
Andreas Keller[7] Monique A.M. Smeets[8] Joel D. Mainland[4,5,*] Sijia Wang[2,9,*]

Manuscript compiled on 4/22/2021

[1] B.L., M.K., and Q.P contributed equally to this work
[2] CAS Key Laboratory of Computational Biology, Shanghai Institute of Nutrition and Health, University of Chinese Academy of Sciences, Chinese Academy of Sciences, China
[3] Department of Skin and Cosmetics Research, Shanghai Skin Disease Hospital, Tongji University School of Medicine, Shanghai, China
[4] Monell Chemical Senses Center, Philadelphia, PA 19104, USA
[5] Department of Neuroscience, University of Pennsylvania, Philadelphia, PA 19104, USA
[6] Unilever Research & Development, Colworth, UK
[7] Laboratory of Neurogenetics and Behavior, The Rockefeller University, New York, NY 10065 USA
[8] Unilever Research & Development, Rotterdam, The Netherlands
[9] Center for Excellence in Animal Evolution and Genetics, Chinese Academy of Sciences, Kunming 650223, China

### Supporting Data

See supporting Files.
**SI Data 1.** Significant Discovery Cohort Associations (p< 5x10-8). Abbreviation: CHR=chromosome; BP=base pair position.
**SI Data 2.** Meta-Analysis Results. Shown are all SNPs that are significantly (p<5x10-8) associated with any tested phenotype in the meta-analysis including both the discovery and validation cohorts. CHR:BP are the chromosome and base-pair coordinates according to human reference genome GRCh37. A1_meta and A2_meta are the two possible nucleotides at each location. The "Direction_meta" column describes the direction of the effect from A1_meta to A2_meta. Both concentrations of odor are included (dilution1 = lower concentration; dilution2 = higher concentration; see main Table 2 for odor concentrations).

### Supporting Tables

**SI Table 1** Phenotype Heritability
**SI Table 2** Frequency of Linked SNPs in OR4D6 Associated with Galaxolide Perception **SI Table 3** Associations Between Galaxolide and SNPs of Other Reported Musk-Related ORs
**SI Table 4** Fine Mapping Analysis

**SI Table 5** OR Haplotypes Tested in the Cell-based Assay for Activation by Galaxolide (OR4D6 Cluster)
**SI Table 6** OR Haplotypes Tested in the Cell-based Assay for Activation by 3M2H (OR51B2 Cluster)

| | Intensity | | Pleasantness | |
|---|---|---|---|---|
| Odor | $h^2$ | SE | $h^2$ | SE |
| β-ionone | 0.38 | 0.31 | 0.00 | 0.31 |
| 3M2H | 0.24 | 0.34 | 0.51 | 0.32 |
| Galaxolide | 0.33 | 0.28 | 0.20 | 0.32 |
| Cis-3-hexenol | 0.00 | 0.25 | 0.00 | 0.28 |
| Decylaldehyde | 0.15 | 0.25 | 0.00 | 0.29 |
| Androstenone | 0.18 | 0.29 | 0.46 | 0.37 |
| Caproic acid | 0.01 | 0.29 | 0.35 | 0.28 |
| Galbanum oxathiane | 0.20 | 0.28 | 0.15 | 0.30 |
| MixB | 0.19 | 0.27 | 0.00 | 0.28 |
| MixA | 0.00 | 0.28 | 0.18 | 0.27 |

Abbreviations: $h^2$=heritability, SE=standard error

**SI Table 1:** Heritability of ranked intensity and ranked pleasantness of 10 odors estimated by GCTA software using LD-pruned variants (143,988 SNPs with $r^2 < 0.2$) from the discovery study.

**Discovery Cohort**

| rs14535412 M263T | rs1453542 S151T: S/S | S/T | T/T |
|---|---|---|---|
| M/M | 582 | 0 | 0 |
| M/T | 4 | 353 | 0 |
| T/T | 0 | 2 | 62 |

n=1003

**Validation Cohort**

| rs14535412 M263T | rs1453542 S151T: S/S | S/T | T/T |
|---|---|---|---|
| M/M | 168 | 0 | 0 |
| M/T | 69 | 83 | 0 |
| T/T | 15 | 5 | 17 |

n=357

**SI Table 2:** Frequency of the two SNPs in OR4D6, rs1453541 (M263T) and rs1453542 (S151T) in discovery and validation cohorts. Haplotypes with the T variant from S151T always have the T variant from M263T.

| Gene | SNP | Variant Allele | p-value | p-value after controlling for top SNP |
|---|---|---|---|---|
| Activated by Muscone | | | | |
| OR5AN1 | rs7941190 | G | $2.98 \times 10^{-8}$ | 0.6811 |
| Broadly-tuned musk receptor | | | | |
| OR5A2 | rs1453547 | A | $4.19 \times 10^{-17}$ | 0.2863 |
| OR5A2 | rs17153691 | G | $3.79 \times 10^{-6}$ | 0.2470 |
| Activated by Nitro musks | | | | |
| OR1A1 | rs4325604 | T | $6.48 \times 10^{-1}$ | 0.8897 |
| OR1A1 | rs769427 | T | $4.37 \times 10^{-1}$ | 0.6239 |
| Discovery Cohort Data; n=1003 | | | | |

**SI Table 3:** The associations between Galaxolide and SNPs of other reported musk-related ORs in the discovery cohort (n=1003) before controlling for the top associated variants (SNPs in OR4D6). OR5AN1 and OR5A2 are in the same LD-band as OR4D6 (see main Figure 2.), meaning variants in these ORs are more likely to be inherited with the SNPs from OR4D6. After performing an additional analysis controlling for the top associated SNPs in OR4D6 (p-value after controlling for top SNP), we found no additional significant signal.

| Odor | CHR | SNP | Pos(hg19) | Gene | Posterior Probability |
|---|---|---|---|---|---|
| β-ionone | 11 | rs6591536 | 59211188 | OR5A1 | 0.50 |
| β-ionone | 11 | rs7941591 | 59211265 | OR5A1 | 0.50 |
| Galaxolide | 11 | rs1453542 | 59224885 | OR4D6 | 0.42 |
| Galaxolide | 11 | rs1453541 | 59225221 | OR4D6 | 0.58 |
| 3M2H | 11 | rs10837814 | 5345128 | OR51B2 | 1.00 |

Abbreviations: CHR=Chromosome, Pos(hg19)=base pair position from genome build hg19

**SI Table 4:** Shown here are all SNPs in the 99% credible set from the fine mapping analysis. For each odor intensity phenotype, we examined SNPs 200kb upstream and downstream from the top associated SNP. We used PAINTOR to calculate posterior probability based on functional annotation linkage disequilibrium. In the case of two highly linked SNPs, such as with OR5A1 and OR4D6, the posterior probabilities sum to 99%.

| OR | Variant | rs # | Explanation |
|---|---|---|---|
| OR4D6 (1) | Reference | | Top association hit in discovery cohort, reference sequence |
| OR4D6 (2) | Consensus | | Top association hit in discovery cohort, consensus sequence |
| OR4D6 (3) | D96G<br>**S151T**<br>**M263T** | rs1453543<br>**rs1453542**<br>**rs1453541** | Top association in the discovery cohort, variant haplotype |
| OR4D6 (4) | M59V<br>D96G<br>**S151T**<br>**M263T** | rs1453544<br>rs1453543<br>**rs1453542**<br>**rs1453541** | Top association in the discovery cohort, variant haplotype |
| OR4D6 (5) | D96G | rs1453543 | Haplotype of OR4D6 not associated with changes from reference |
| OR5A1 (1) | Reference | | Top association in validation cohort, reference haplotype |
| OR5A1 (2) | D183N | rs6591536 | Top association in validation cohort, variant haplotype |
| OR5AN1 (1) | Reference | | In OR4D6 cluster |
| OR5A2 (1) | Reference | | In OR4D6 cluster |
| OR5A2 (2) | P172L | rs1453547 | Variant of OR5A2 in OR4D6 cluster |

**SI Table 5:** Olfactory receptor haplotypes (hg19) tested in the cell-based assay for activation by Galaxolide (OR4D6 Cluster). The bolded variants are the SNPs associated with change in Galaxolide perception. OR4D6 (2) is a consensus version of OR4D6 across 10 closely related species (Trimmer et al., 2019 (10); Ikegami et al., 2020 (30))

| OR | Variant | rs # | Explanation |
|---|---|---|---|
| OR51B2 | Reference | | Top association hit in validation cohort, reference sequence |
| OR51B4 | V36I<br>M147T | rs7118113<br>rs10837771 | In OR51B2 cluster, and nearest OR to top association in discovery study (SNP in non-coding region) |
| OR52A1 | Reference | | The top association in discovery study (SNP in a non-coding region) is an eQTL affecting expression of OR52A1, making OR52A1 the putative responding receptor |
| OR52A4 | Reference | | In OR52A1 cluster |
| OR52A5 | Reference | | In OR52A1 cluster |
| OR51B6 | Reference | | In OR51B2/4 cluster |
| OR51B5 (1) | Reference | | In OR51B2/4 cluster |
| OR51B5 (2) | I102T<br>P160L | rs11036912<br>rs4910551 | Variant of OR51B5 in the OR51B2/4 cluster |

**SI Table 6:** Olfactory receptor haplotypes tested in cell assay for activation by 3M2H (OR51B2/4 Cluster).

### Supporting Figures

**SI Fig.1** Phenotype Distribution
**SI Fig.2** 3M2H/OR51B1 LD-band Cell-based Assay Results
**SI Fig.3** Androstenone Intensity by OR7D4 RT/WM genotype in Discovery Cohort
**SI Fig.4** Natural Selection Results
**SI Fig.5** PCA of Population Structure for Discovery, Validation, and 1000 Genomes Data

A)

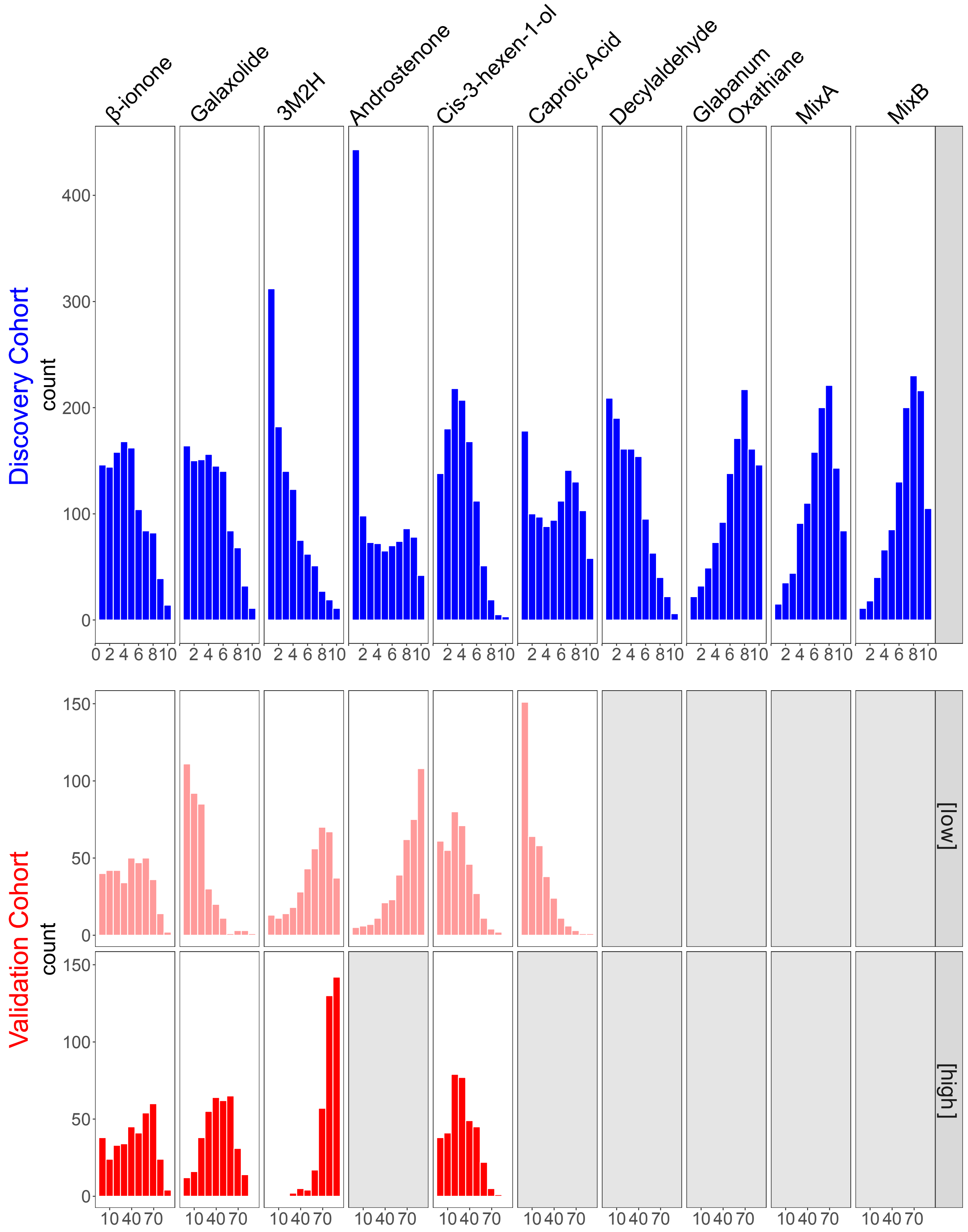

β-ionone
Galaxolide
3M2H
Androstenone
Cis-3-hexen-1-ol
Caproic Acid
Decylaldehyde
Glabanum Oxathiane
MixA
MixB
Discovery Cohort
count
400
300
200
100
0
0 2 4 6 8 10
2 4 6 8 10
Validation Cohort
count
150
100
50
0
10 40 70
[low]
[high]

**B)**

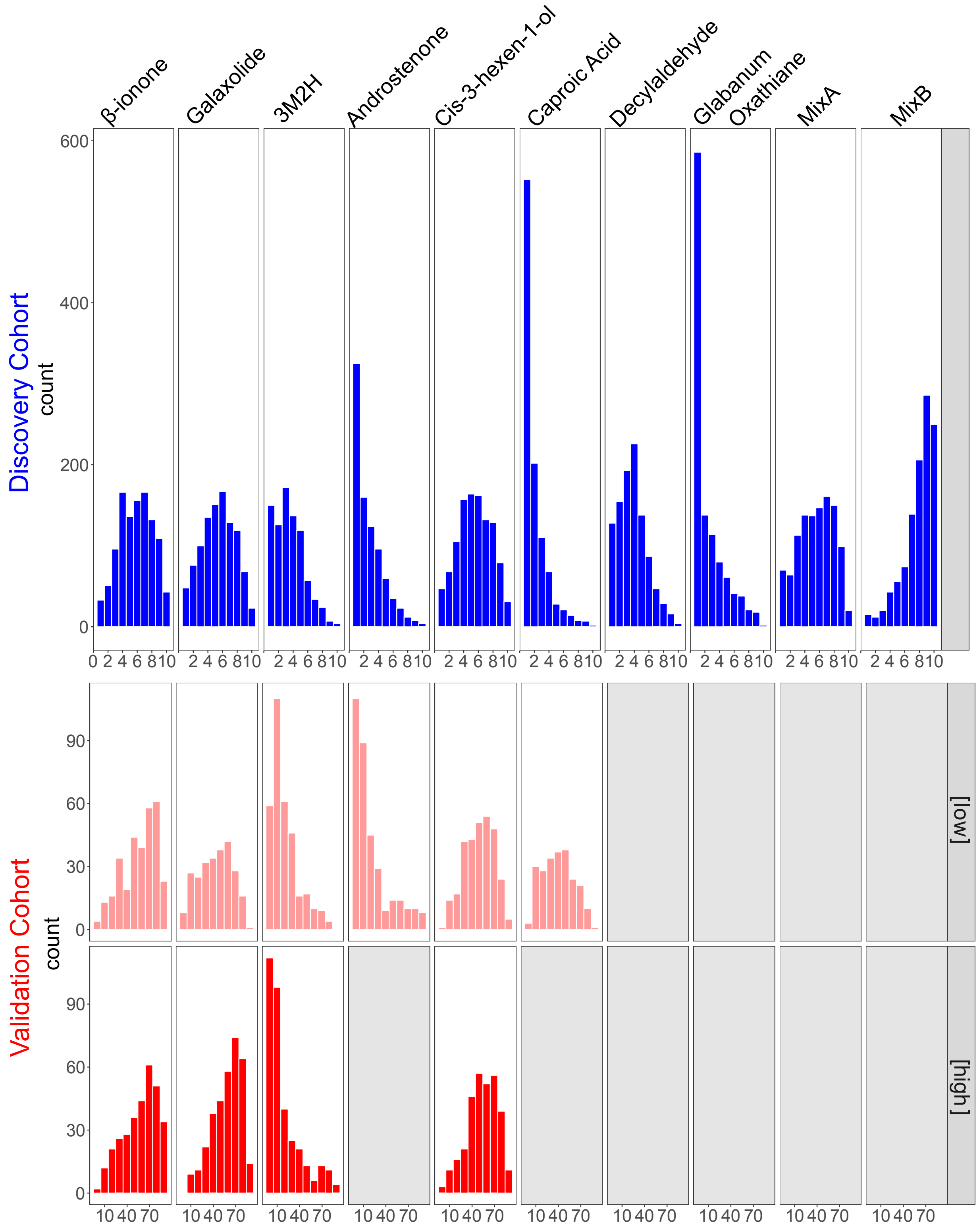


**SI Figure 1:** Distribution of ranked intensity (A) and pleasantness (B) ratings for odors in the discovery (blue) and replication (red) studies. A grey box indicates the phenotype was not tested.

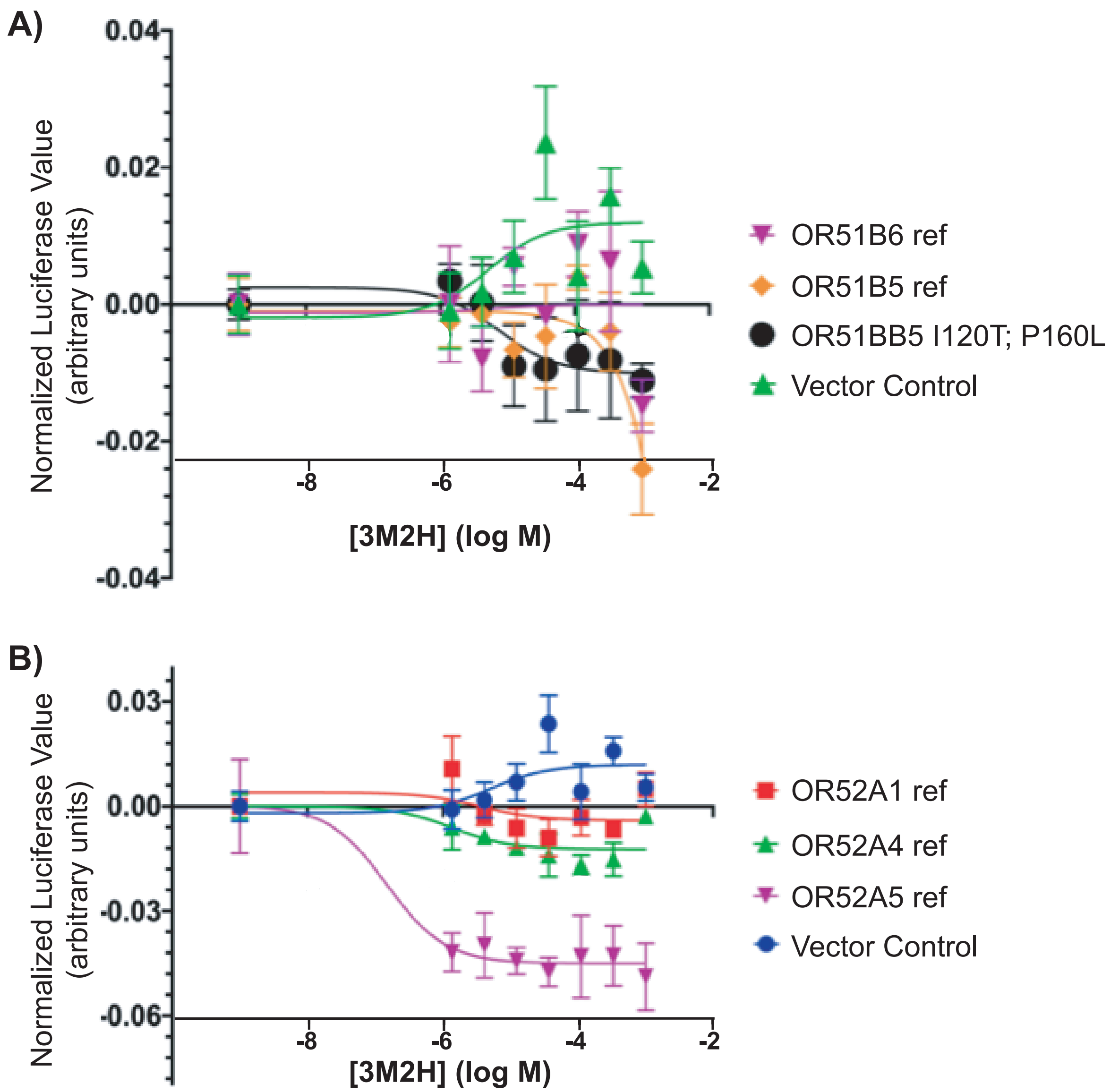


**SI Figure 2:** Cell-based assay results for 3M2H against other receptors in the A) OR51B2 and B) OR52A1 clusters. No receptors responded significantly above the vector control (Rho). Luciferase values were normalized by RL readings and then baselined to zero by subtracting the response of the no-odor control.

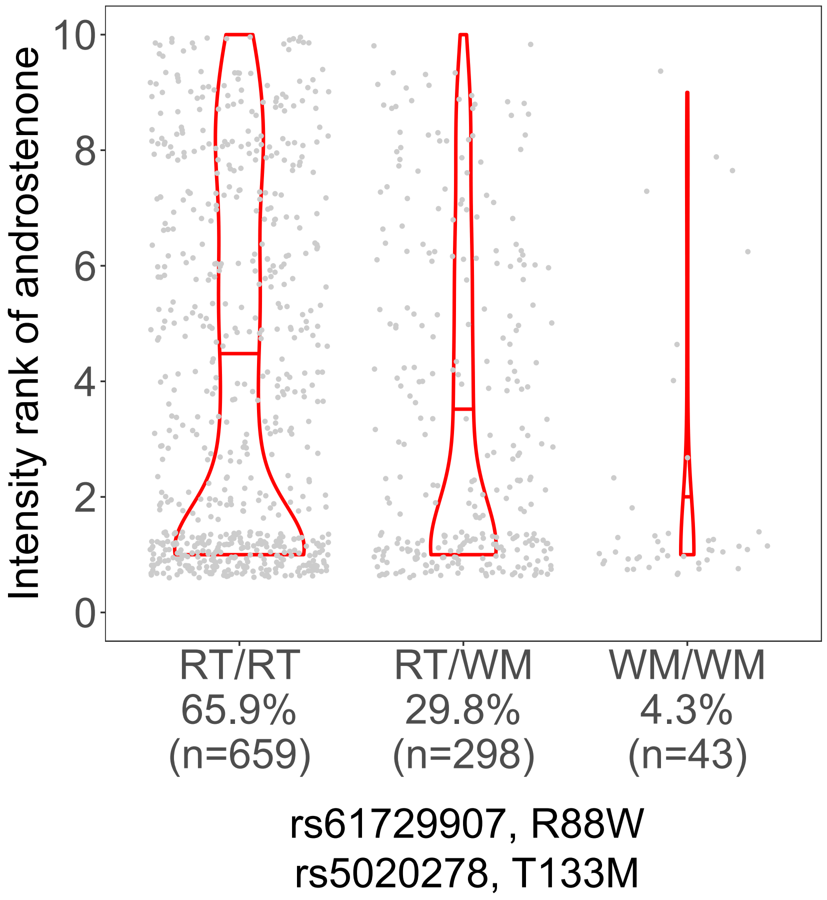


**SI Figure 3:** Intensity perception of androstenone is associated with RT/WM haplotype of OR7D4 in the discovery cohort.

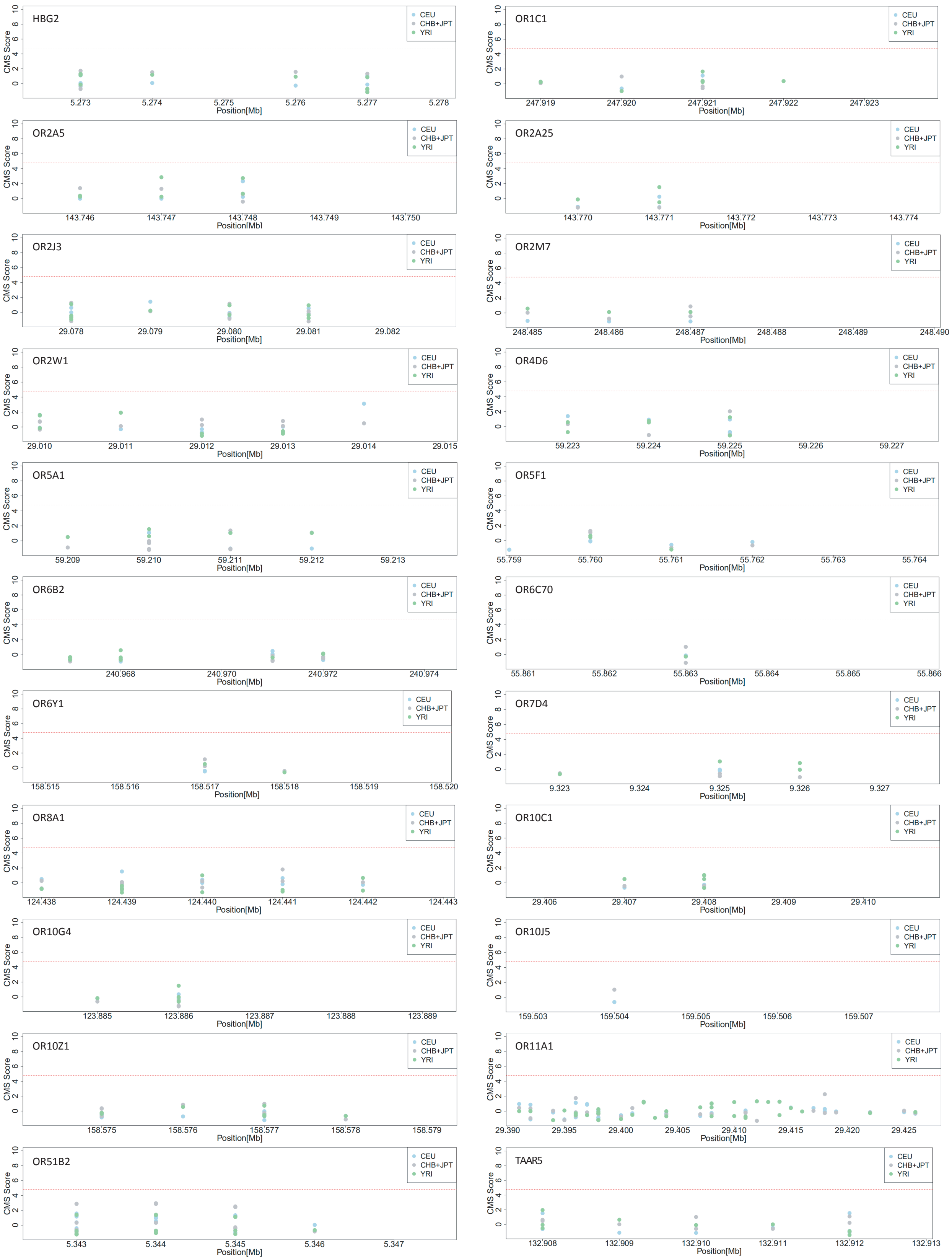


**SI Figure 4:** Results for natural selection on candidate OR gene regions (±2kb). CMS scores are plotted against chromosome position in CEU, CHB+JPT, and YRI populations, shown in blue, gray, and green, respectively. The red dotted line represents the significance threshold (top 0.1% CMS score: 4.791). No enrichment for high CMS scores (top 0.1%) is found within the genes, indicating the examined SNPs are not subject to natural selection.

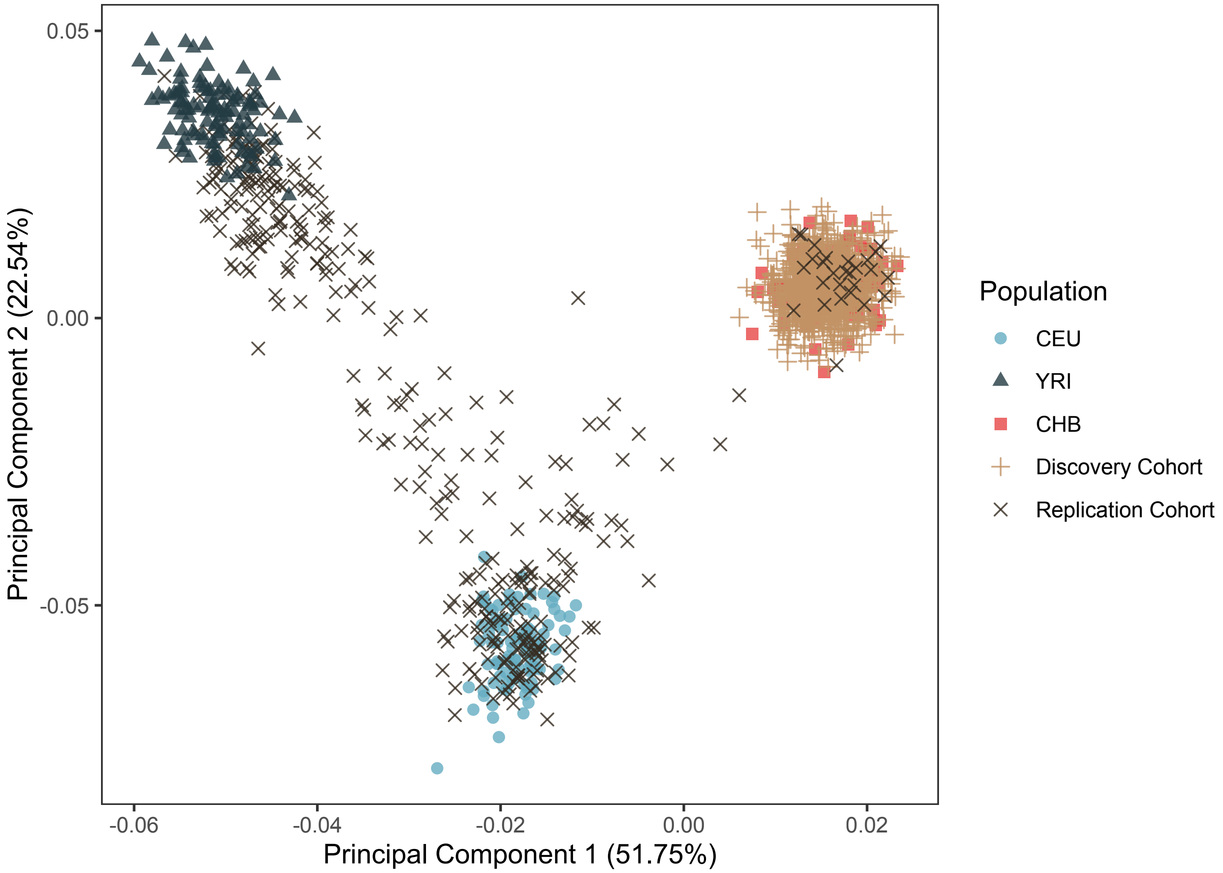


**SI Figure 5:** Population structure analysis reveals relative homogeneity of the discovery population (tan +) compared to the validation population (black x) ($p<2.2x10^{-16}$). Shown are the first two principle components calculated from all variants genotyped in both the discovery and validation cohorts. Representative populations from the 1000 Genomes Project: Han Chinese in Beijing (CHB, n=97; red □), Utah residents with Northern and Western European ancestry from the CEPH collection (CEU, n=86; blue ◯), and Yoruba in Ibadan, Nigeria (YRI, n=88; black △) are plotted for context. The discovery population overlapped with the CHB population (mean distance to CHB=0.001, CEU=0.07, YRI=0.07).
